# Impact of K-Pg Mass Extinction Event on Crocodylomorpha Inferred from Phylogeny of Extinct and Extant Taxa

**DOI:** 10.1101/2021.01.14.426715

**Authors:** Andrew F. Magee, Sebastian Höhna

## Abstract

Crocodilians and their allies have survived several mass extinction events. However, the impact of the K-Pg mass extinction event on crocodylomorphs is considered as minor or non-existent although other clades of archosaurs, e.g., non-avian dinosaurs and pterosaurs, went extinct completely. Previous approaches using fossil occurrence data alone have proven inconclusive. In this paper, we take a phylogenetic approach using extant and extinct species. The time-calibrated phylogeny of extant crocodilians provides insights into the pattern of recent biodiversity changes whereas fossil occurrence data provide insights about the more ancient past. The two data sources combined into a single phylogeny with extinct and extant taxa provide a holistic view of the historical biodiversity. To utilize this combined data and to infer the impact of the K-Pg mass extinction event, we derive the likelihood function for a time-varying (episodic) serially sampled birth-death model that additionally incorporates mass extinctions and bursts of births. We implemented the likelihood function in a Bayesian framework with recently developed smoothing priors to accommodate for both abrupt and gradual changes in speciation, extinction and fossilization rates. Contrary to previous research, we find strong evidence for the K-Pg extinction event in crocodiles and their allies. This signal is robust to uncertainty in the phylogeny and the prior on the mass extinctions. Through simulated data analyses, we show that there is high power to detect this mass extinction and little risk of false positives.

## Introduction

Crocodylomorpha (living crocodilians and their extinct relatives) is an ancient clade that first appeared in the late Triassic (Markwick 1998; Brochu 2003). Crocodylomorphs have survived several mass extinction events, including the Triassic-Jurassic extinction event (201.3 Mya), the Jurassic-Cretaceous extinction (145 Mya), the Cretaceous-Paleogene extinction event (K-Pg extinction, 66 Mya) and the Eocence-Oligocene extinction event (33.9 Mya). The fossil records shows significantly higher historical diversity with a peak in Jurassic compared to the diversity of extant crocodilians (Bronzati et al. 2015). Previous studies have found evidence for the Triassic-Jurassic extinction (Markwick 1998) and the Jurassic-Cretaceous extinction (Tennant et al. 2016) although some debate of the true impact remains (Fanti et al. 2016). Most surprisingly, all previous studies assessed the impact of the K-Pg mass extinction as minor or non-existent (Markwick 1998; Brochu 2003; Bronzati et al. 2015; Mannion et al. 2015) although sister clades, such as non-avian dinosaurs and pterosaurs, with similar lifestyle (e.g., body size, habitat and food sources) and anatomy where severely impacted and went extinct 66 Mya.

The fossil record alone might not be sufficient to answer what impact the K-Pg mass extinction on crocodylomorphs had. Alternatively, time-calibrated phylogenies have been used successfully to infer the impact of mass extinction events (Crisp and Cook 2009; May et al. 2016; Culshaw et al. 2019), e.g., on fishes (Tetraodontiformes) (Arcila and Tyler 2017) and geckos (Brennan and Oliver 2017). However, the signal of mass extinction erodes the further back the event was in the past (May et al. 2016). The age of the crown group of living crocodylomorphs (Crocodylia) dates back to the Campanian with estimates ranging between 90 to 118 Mya (Oaks 2011; Wilberg et al. 2019). Even considering the older estimates of the age of the crown group, only three lineages that crossed the K-Pg boundary have living descendants (ancestral alligators, crocodiles, and gharials). Thus, a time-calibrated phylogeny of living crocodilians alone is also not sufficient to show the impact of the K-PG mass extinction on Crocodylomorphs.

Here, we develop a new statistical approach using phylogenies of both living and extinct species. Our approach extends previous fossilized-birth-death models (Stadler 2010; Heath et al. 2014; Gavryushkina et al. 2014). Specifically, our new model includes piecewise-constant (episodic) rates of speciation, extinction and fossilization and instantaneous events of mass extinction with an estimated survival probability (Höhna 2015; May et al. 2016). For mathematical completeness and availability for future research, we also complement the continuous events of speciation and fossilization with instantaneous events of bursts (e.g., Oaks 2019) and mass fossilization. We also draw a link to infectious disease modeling and present our birth-death-sampling process as a generalization of both macroevolutionary and phylodynamic models (Stadler et al. 2013; Kühnert et al. 2014).

We demonstrate our new model on a recently published phylogeny of crocodylomorphs including extinct and extant species (Wilberg et al. 2019). We use a reversible-jump Markov chain Monte Carlo algorithm (Green 1995) to infer whether a mass extinction event has impact our study group or not. We use Horseshoe Markov random field (HSMRF) smoothing priors (Magee et al. 2020) on speciation, extinction and fossilization rates to allow for abrupt changes while favoring constant rates over rate variation (Occam’s razor principle). We infer an overwhelming support for a mass extinction event about 68 Mya.

## Results

### The Generalized Episodic Fossilized-Birth-Death Process

We derived the probability density function of a phylogeny with extant and extinct tips under the generalized episodic fossilized-birth-death process (see Equation 1 in our *Methods*). Our generalized model combines previous special cases for macroevolution (Stadler 2011; Gavryushkina et al. 2014; Höhna 2015; May et al. 2016) and phylodynamic models (Stadler et al. 2013; Gavryushkina et al. 2014) and extends the existing models to include mass extinction events as well as bursts of birth events. Our generalized episodic fossilized-birth-death process collapses to exactly the same processes and probability density function of these special cases (see Supporting Information). Thus, our model provides a flexible framework to explore new and previously described models within the same framework. Most importantly, our model enables robust estimation of variation in diversification rates using phylogenies with extant and/or extinct taxa. The probability density function is implemented in the software `RevBayes` (Höhna et al. 2016). This implementation allows both for flexible Bayesian parameter estimation and use of the generalized episodic fossilized-birth-death process as a prior distribution for divergence time estimation (e.g., see Heath et al. 2014).

### K-Pg mass extinction in Crocodylomorpha

We tested for the effect of mass extinctions in Crocodylomorpha using the dataset of Wilberg et al. Wilberg et al. (2019). The tree is 243.5 million years old, and includes 14 of the 23 known extant species and 128 fossil-tips (Wilberg et al. 2019; Oaks 2011). To account for the possible effect of background rate variation, and for the possibility that a mass extinction is followed by a rapid burst of speciation (Crisp and Cook 2009), we employed time-varying priors on the speciation, extinction, and sampling rates through time (Magee et al. 2020). We computed Bayes factors to test for the signal of a mass extinction event anytime along the phylogeny. We explored the robustness of results using six published phylogenies by Wilberg et al. (Wilberg et al. 2019) and five different priors on the expected number of mass extinction events.

We find very strong evidence (2ln Bayes Factor > 10 for all dataset and prior combinations) for a mass extinction event approximately 68 Mya, which is most likely a signal of the K-Pg mass extinction (Figure 1). We find weak positive evidence for a signal of the Tr-J mass extinction, though the timing is less certain. The weaker signal for this ancient mass extinction event is most probably due to the age of the mass extinction event, as the signal of a mass extinction event within the first 25% of the age of the phylogeny is often erased (May et al. 2016). We also find a very weak signal for the J-C mass extinction. Overall, we recovered some signature for all four major mass extinctions that crocodylomorphs have survived and our method shows very strong support for no other mass extinction between the known events. Interestingly, there is a very weak signature of a mass extinction towards the present. This indicates that living crocodilians are in decline and might face yet another mass extinction, which is further corroborated by our estimated speciation and extinction rates (Figure 2).

**Figure 1:**
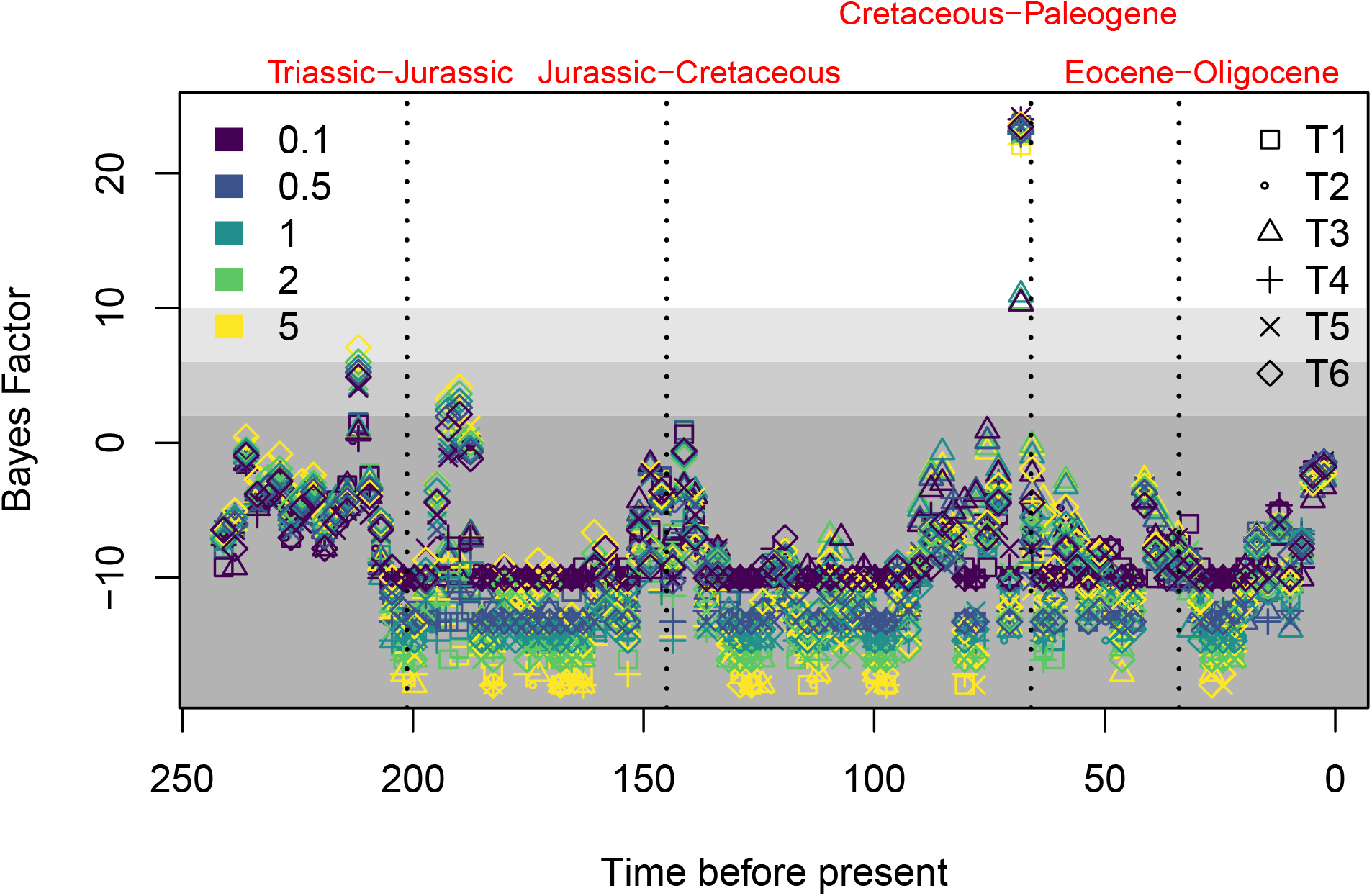
Per-interval support for mass extinctions in the Crocodylomorpha dataset for all 30 combinations of dataset and mass extinction prior. Mass extinctions are allowed only at the end of each interval, and all intervals are fixed. Different mass extinction priors are shown by color, and different datasets by symbols. Shaded regions denote 2ln Bayes Factor cutoffs of 2, 6, and 10, which correspond to weak support, support, and strong support for a mass extinction. Across all analyses there is a strongly supported mass extinction that corresponds to the time of the K-Pg.

**Figure 2:**
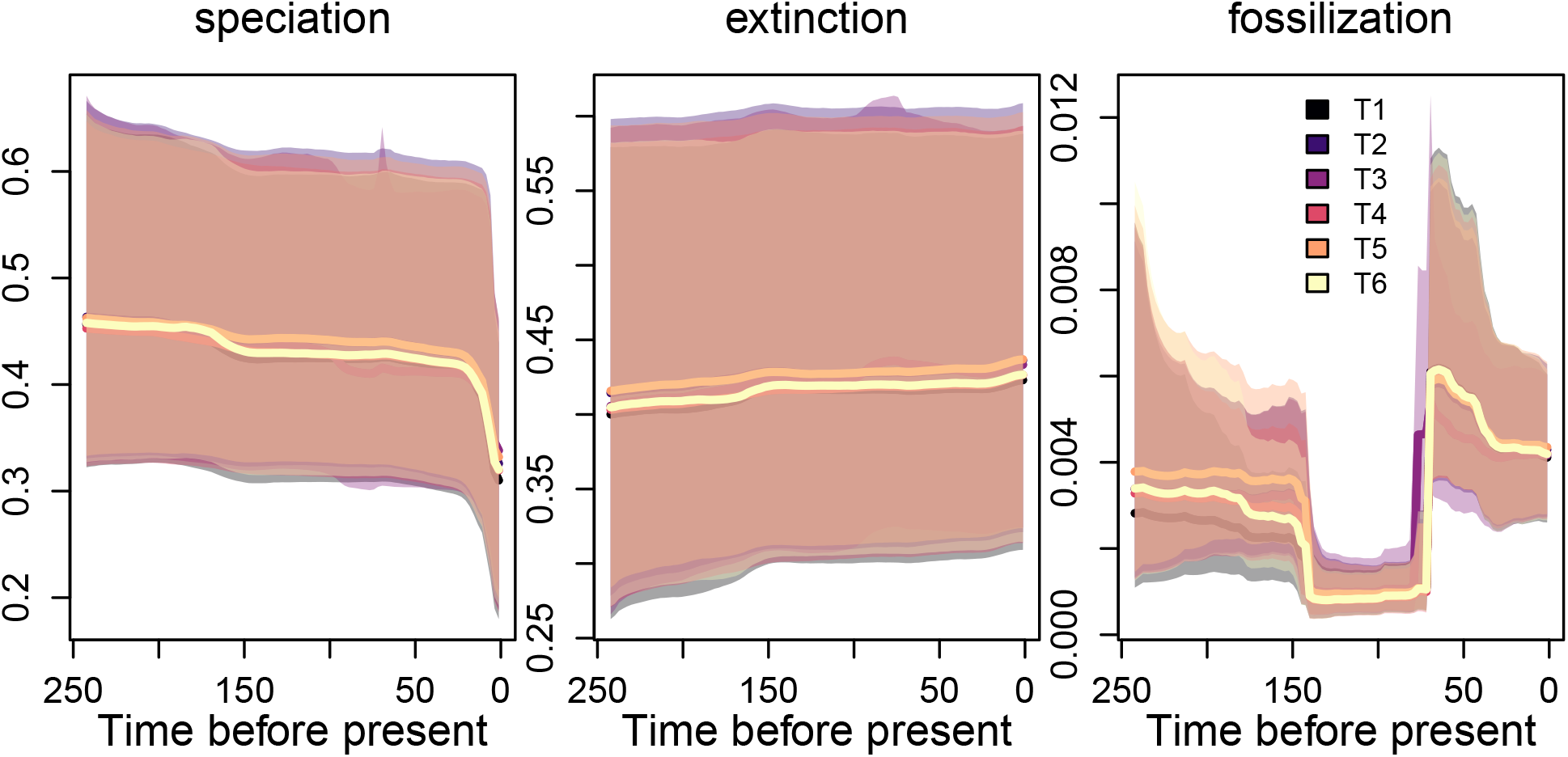
The rates of speciation, extinction, and fossilization through time for one mass extinction prior 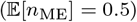 and all datasets. Speciation rates began to decline steeply approximately 25 Mya, leading to a negative rate of net diversification. Estimates are broadly similar across all datasets and mass extinction priors, full plots are available in Figure S1.

Approximately 25 Mya, the speciation rates began a sharp decline, with extinction rates exceeding speciation rates in the recent past (Figure 2). The estimated fossilization rates exhibit substantial variation. However, we emphasize that the estimated fossilization rate is distinct from the actual fossil prevalence at any time. The fossilization rate is a per-lineage rate which depends on the number of lineages in the tree, and its estimate is confounded by fossil recovery and fossil sampling for the specific study. Reassuringly, the estimated speciation, extinction and fossilization rates are in strong agreement regardless of the specific phylogeny and prior probability on the number of mass extinction events (Figure S1).

### Power and False Positive Rate

We explored the power and false-discovery rate of detecting mass extinction events using simulations. We find that our method has a high power to detect mass extinctions, as the simulated K-Pg mass extinction is detected in 82% of simulations (Figure 3). We also find that our method has a low false positive rate, as 4% of simulations with mass extinctions detect a mass extinction at an incorrect time, and 4% of simulations without mass extinctions detect any mass extinction (Figure 3). This simulation shows that our inferred signal of the K-Pg mass extinction is indeed a true signal.

**Figure 3:**
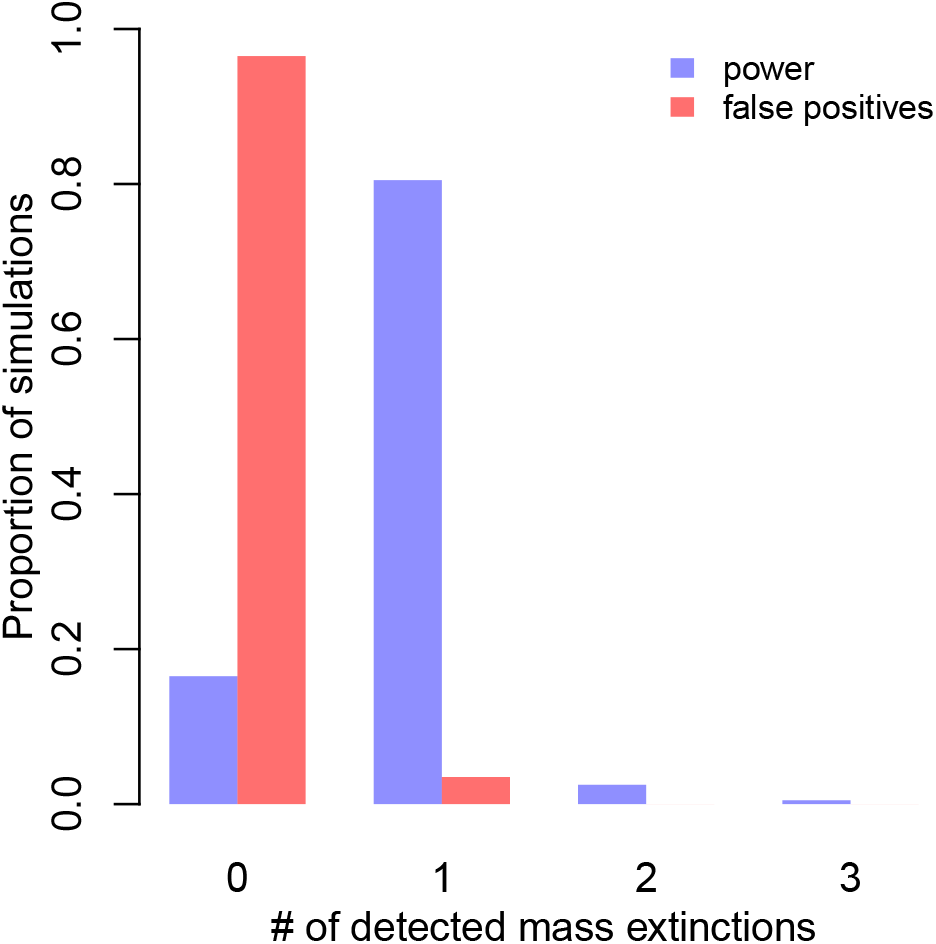
The number of detected mass extinctions across our 400 simulated data analyses. Detection is defined as a mass extinction for which the 2ln Bayes Factor is at least 10. In most simulations without mass extinctions (red), no mass extinctions are inferred, while in most simulations with mass extinctions (blue), one mass extinction is inferred.

### Model Adequacy

Our empirical results provided very strong evidence for an impact of the K-Pg mass extinction on crocodilians. Nevertheless, even the best supported model might be inadequate for a given dataset. We tested model adequacy of both a model with and without the K-Pg mass extinction using posterior predictive simulations. Our results show that both models are equally adequate and inadequate (see supplementary figure S8). Overall, our generalized episodic fossilized birth-death model with mass extinction showed a very good fit except for three summary statistics: Colless’ imbalance statistic (Colless 1982), Tippyness (Rohlf et al. 1990) and the number of sampled ancestors. The deviation from the expected tree balance shows that the observed crocodilian phylogeny was most likely produced by a process of lineage heterogeneous speciation and extinction rates. The empirical number of sampled ancestors, which is zero, is most likely an artifact of the tree construction method which did not allow for fossils to be sampled ancestors (Wilberg et al. 2019). Our model can be modified to enforce zero sampled ancestors by assuming that a fossilization event is always combined with an extinction event. Our results (supplementary material figure S4) show that the inference of the K-Pg mass extinction is robust to the systematic bias of not allowing for sampled ancestors.

### Signal from Extant vs Extinct Species

Previous research debated whether molecular phylogenies from extant taxa need the fossil record to reliably infer historical diversification patterns (Quental and Marshall 2010; Morlon et al. 2011). Our generalized episodic fossilized birth-death process bridges analyses from the fossil record and molecular phylogenies. Nevertheless, we performed analyses using only the extant tips (pruning all fossils) and only using extinct tips (pruning all extant taxa). In the case of crocodilian diversification, the strongest signal is contained in the fossil taxa (see supplementary material figure S2). Thus, molecular phylogenies may indeed need the fossil record to obtain a complete picture of historical diversification processes. However, the importance of the fossil taxa for our analyses is not surprising because the phylogeny by Wilberg et al. contains ten times more fossil taxa than extant taxa and the crown age of the extant taxa is too recent to show a signal of the K-Pg mass extinction or other older mass extinctions.

## Discussion

In this paper, we have described the generalized episodic fossilized birth-death model, which includes three previously ignored parameters and constitutes the most parameter-rich time-dependent birth-death model to date. The new parameters **Λ** (models tree-wide bursts of births), and ***M*** (models tree-wide bursts of deaths, i.e., mass extinctions) will primarily be of interest in macroevolutionary studies, though they may be of use to phylodynamic situations for modeling superspreader events and culling. The new parameter ***R*** (models different treatment probabilities between *ϕ*-sampling and Φ-sampling) will primarily be of interest in phylodynamic applications. The generalized episodic fossilized birth-death model is implemented in `RevBayes` (Höhna et al. 2016), and is therefore available for inferring diversification parameters on fixed trees and for use in full Bayesian joint inference of divergence times and diversification rates.

We applied the generalized episodic fossilized birth-death model to a dataset of 142 crocodylomorphs. In a Bayesian analysis, we find strong evidence for the K-Pg mass extinction in crocodylomorphs, a signal which we show is robust to phylogenetic uncertainty, the prior on mass extinctions, and the exclusion of extant taxa. Using post-hoc analyses of simulated data, we show that we have good power to detect this signal and a low risk of false positives. Further post-hoc analyses of the crocodylomorph dataset show that the primary signal for the K-Pg mass extinction is found in the fossil taxa. In general, we expect that the use of fossil samples should greatly increase the power of phylogenetic birth-death models to detect mass extinctions. An extant tip only contains information about mass extinctions through the branching time that it adds to the tree. Sampled ancestors contain information only through the sampling time. Fossil tips, on the other hand, contain both sources of information (branching time and sampling time) and their inclusion should prove useful to the detection of mass extinctions and variation in diversification rates. Our results do show that diversification-rate estimates change when extant tips are excluded, and thus we advocate for using the largest possible dataset when inferring diversification rates through time and/or mass extinctions.

### The Generalized Episodic Fossilized-Birth-Death Process Notation

We begin with the notation of the generalized fossilized-birth-death process (or birth-death-sampling-treatment process as it might be called in the phylodyamics literature). At any point in time, lineages may speciate (infect another individual), go extinct (become noninfectious), or fossilize (be sampled). Additionally, to allow for phylodynamic applications, when a lineage is sampled it may be treated and then become noninfectious (go extinct). Each of these four types of events may occur continuously over time affecting a single lineage (Figure 4 a-d) or instantaneously at a pre-defined time affecting all lineage alive at the time (Figure 4 e-h). The continuously occurring events are governed by time-varying rates λ(*t*), *μ*(*t*), *ϕ*(*t*), and *r*(*t*) for speciation, extinction, fossilization (or sampling) and treatment. The instantaneous, tree wide events are defined by a vector of pairs of times and probabilities (Λ_*i*_, *M_i_*, Φ_*i*_, and *R_i_* respectively). For a summary of the notation and model parameters, see Table 1.

**Figure 4:**
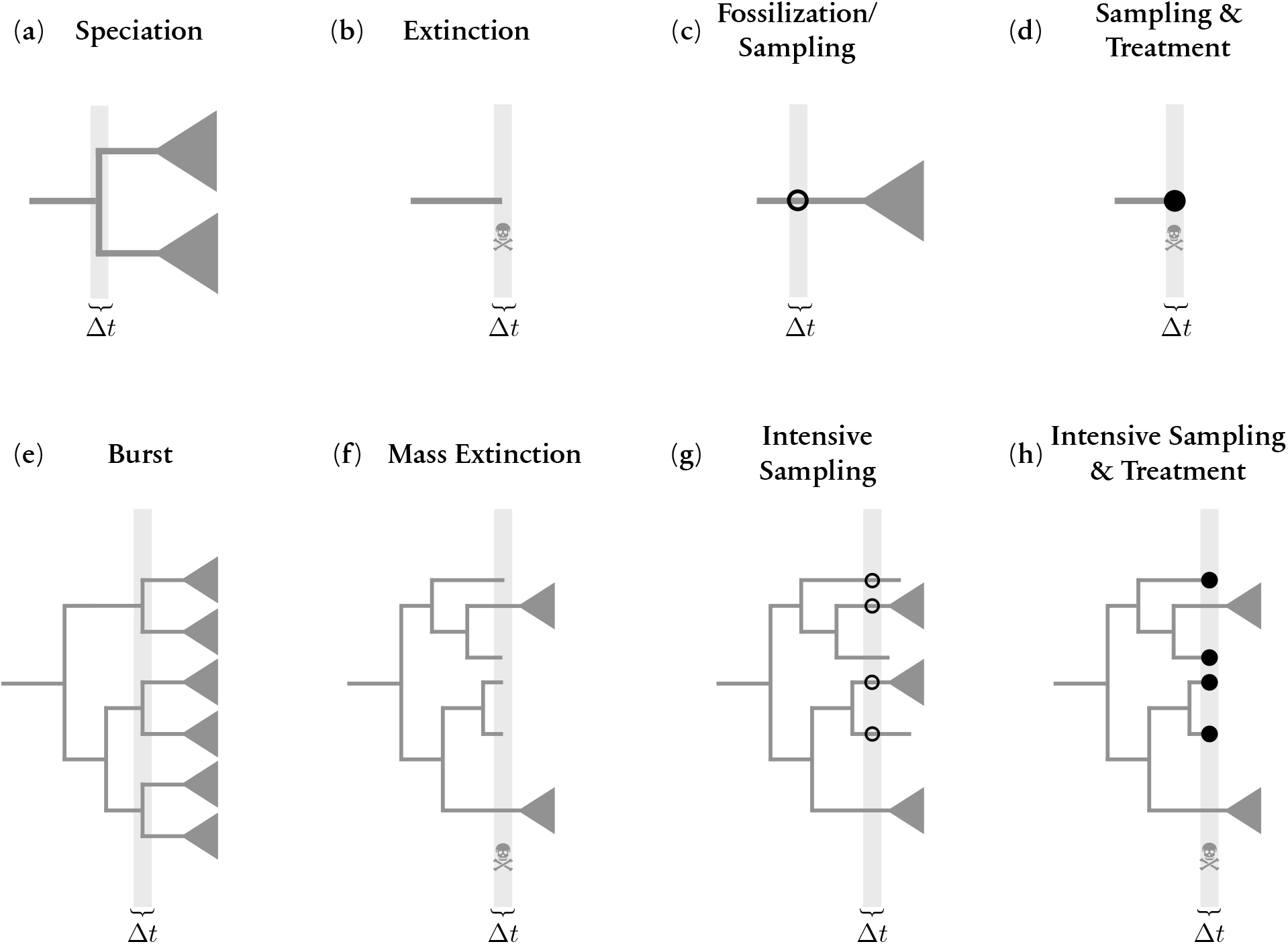
Schematic of possible events. The top row shows continuously occurring events affecting a single lineage and the bottom row shows the same types of events but instantaneously and affecting all lineages (tree wide). The types of events are speciation, extinction, fossilization or sampling, and sampling with treatment.

**Table 1:** Model parameters and their interpretation

| Parameter | Interpretation |
| --- | --- |
| $t_{or}$ | the starting time of the process (can be the origin or the most recent common ancestor). |
| $\lambda(t)$ | *birth* rate at time $t$. |
| $\mu(t)$ | *death* rate at time $t$. |
| $\phi(t)$ | (serial) *sampling* or *fossilization* rate at time $t$. |
| $r(t)$ | probability that a ϕ-sampling event at time $t$ causes a death event. |
| $l$ | the number of breaks between intervals/episodes. |
| $s$ | the beginning times of the episodes/intervals, $s_0, \dots, s_l$, with $s_0 = 0$. |
| $\Lambda$ | vector of $l$ burst probabilities. At time $s_i$ ($i \geq 1$) every lineage *speciates* with probability $\Lambda_i$. |
| $M$ | vector of $l$ extinction probabilities. At time $s_i$ ($i \geq 1$) every lineage *dies* with probability $M_i$. |
| $\Phi$ | vector of $l + 1$ sampling probabilities. At time $s_i$ every lineage is *sampled* with probability $\Phi_i$. |
| $R$ | vector of $l$ treatment probabilities. At time $s_i$ ($i \geq 1$) every sampled lineage dies with probability $R_i$. |
| $\mathcal{A}$ | set of ages of all ϕ-sampled *sampled ancestors*. |
| $\mathcal{B}$ | set of all branch segments, such that $t_o$ is the beginning of the segment and $t_y$ the end ($t_y < t_o$). |
| $\mathcal{F}$ | set of ages of all ϕ-sampled *tips*. |
| $\mathcal{N}$ | set of ages of all non-burst bifurcation (birth) events. |
| $L(t)$ | the number of lineages in the reconstructed tree alive at time $t$. |
| $H$ | total number of ϕ-samples. |
| $I_i$ | number of Φ-samples at the $i$th Φ-sampling event. |
| $I$ | total number of Φ-samples, with $I = \sum_{i=0}^l I_i$. |
| $T_i$ | number of tips with sampled descendants in a given Φ-sampling event. |
| $K_i$ | number of lineages with a burst event at the end of episode $i$. |
| $K$ | total number of $K = \sum_{i=0}^l K_i$. |

### The probability density of a phylogenetic tree

Our derivation of the probability density of a phylogenetic tree with extinct and extant taxa under the generalized episodic fossilized-birth-death process breaks up each branch of the tree into segments belonging only to one epoch (Figure 5). This derivation follows the same logic of Stadler et al. Stadler et al. (2013) and Gavryushkina et al. Gavryushkina et al. (2014) but additionally includes mass extinction and burst events. The probability density of a phylogenetic tree Ψ is

**Figure 5:**
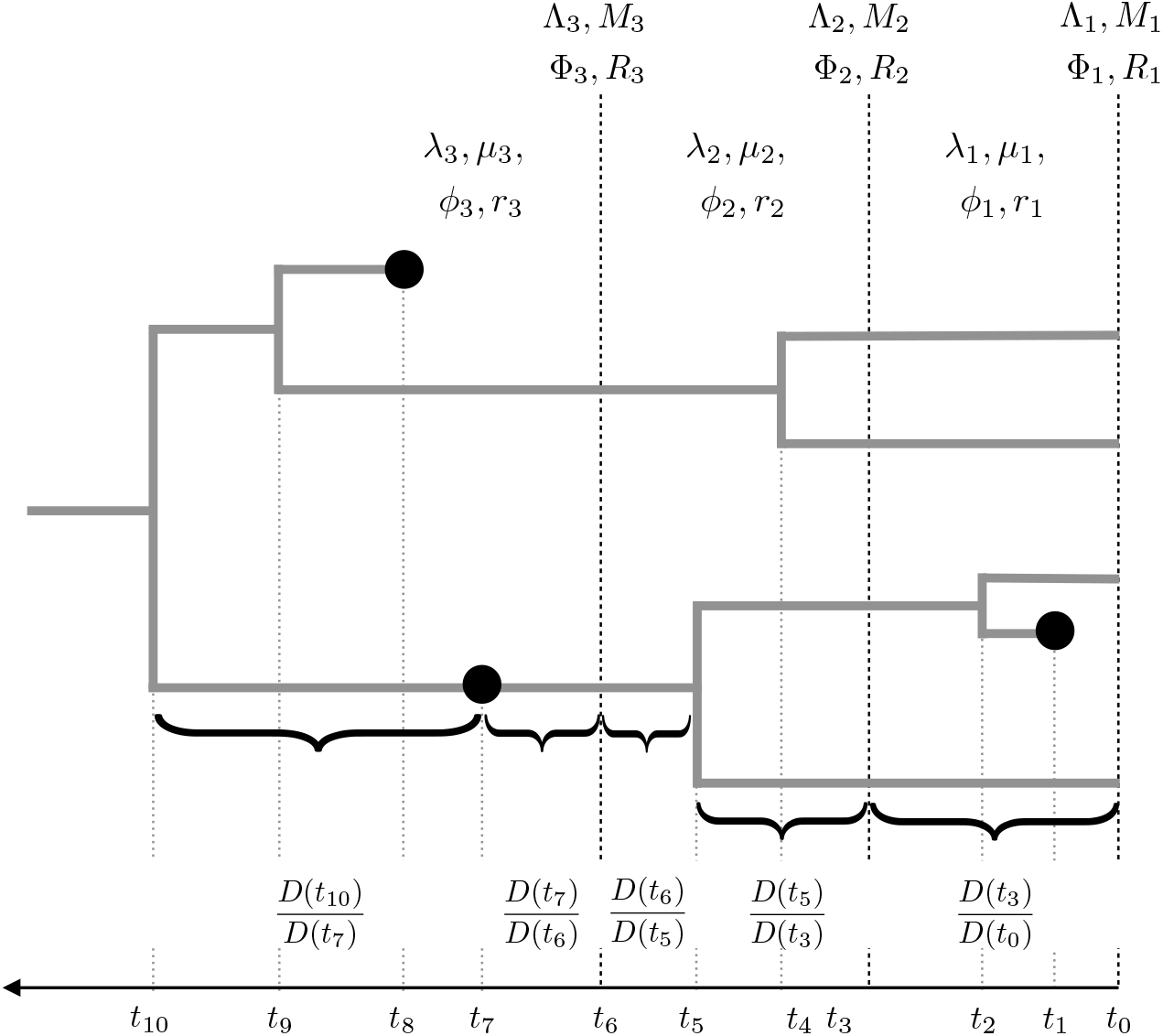
Schematic decomposition of probability density function for our generalized episodic fossilized birth-death process. In this example, we have three epochs and thus three rates (e.g., {λ_1_, λ_2_, λ_3_}) for each continuous type event and three probabilities (e.g., {Λ_1_, Λ_2_, Λ_3_}) for each instantaneous tree-wide event. Each branch is broken into branch segments that starts (tipwards) at *t_y_* and ends at *t_o_* so that within each branch segment no epoch switch, speciation or fossilization event occurs. We compute the probability of observing each branch segment by 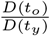, which constitutes the core part of the probability density computation.

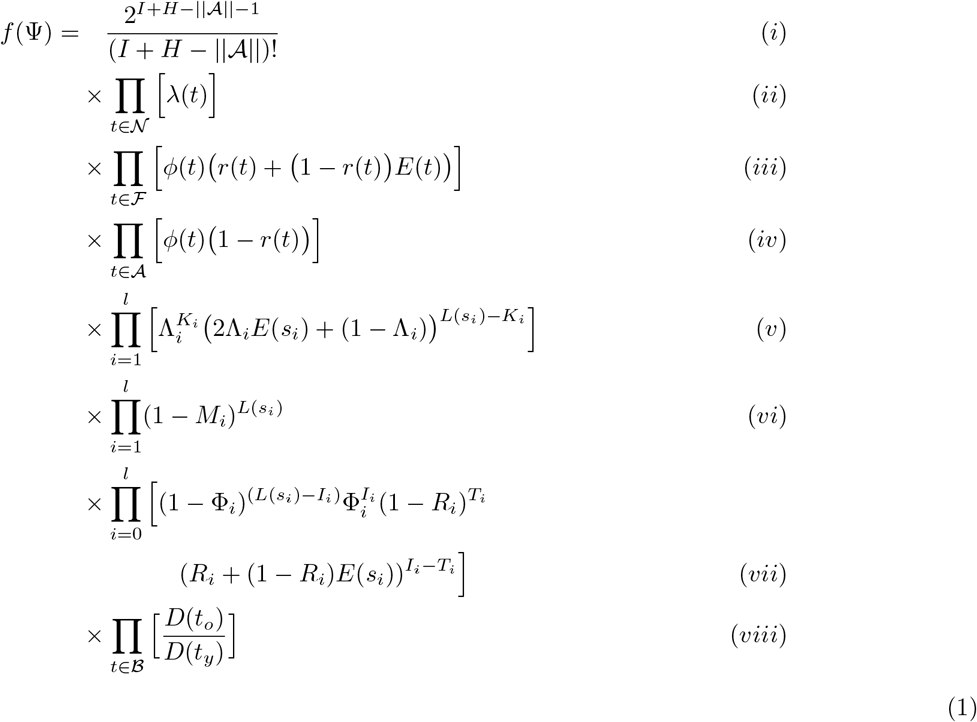

where (i) the probability of the topology, (ii) the probability of the observed serial speciation events, (iii) the probability of the serially-sampled (fossil) tips (Figure 4c-d), (iv) the probability of the sampled ancestors (fossils) (Figure 4c), (v) the probability of the observed and unobserved speciation events at tree-wide speciation burst events (Figure 4e), (vi) the probability of all lineages surviving a tree-wide mass extinctions events (Figure 4f), (vii) the probability of all the observed sampling times at given tree-wide sampling events (Figure 4g), (viii) the probability of the observed branch segments (Figure 5).

### Probability of no sampled descendant E(t)

We begin our derivation by tracking *E*(*t*), the probability that a lineage alive at time *t* has no sampled descendants when the process is stopped. Let us define the differential equation for *E* in a small time interval Δ*t* to be,

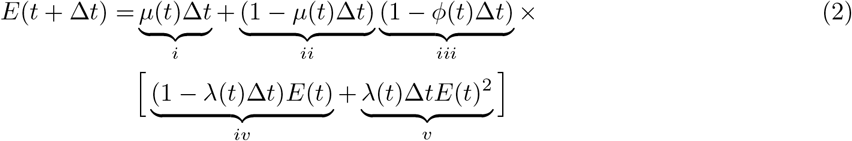

which accounts for the three scenarios that can occur during the interval Δ*t*, (1) the lineage dies (*i*), (2) no death (*ii*) and no serial sampling (*iii*) and no birth (*iv*), (3) no death (*ii*) and no serial sampling (*iii*) and a birth event (*iv*). Cases (2) and (3) require the extant lineage(s) to eventually die or remain unsampled, hence *E*(*t*) and *E*(*t*)^2^. Rearranging the equation, discarding all terms on the order of Δ*t*^2^, and dividing by Δ*t*, we get,

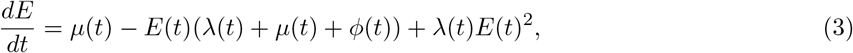

where again we can see three cases, (1) the process dies, (2) nothing happens followed by *E*(*t*), and (3) a birth followed by both lineages not being observed, *E*(*t*)^2^.

We now make the assumption that λ(*t*), *μ*(*t*), and *ϕ*(*t*) are piecewise-constant (see Stadler 2011; Höhna 2015). Additionally, we assume that these rates share the same intervals, i.e., changes in rates may occur at the same time points. We keep track of these times with the vector *s*, of which *l* elements must be specified, *s*_1_,…, *s_l_*. We implicitly specify *s*_0_ = 0 and *s*_*l*+1_ = ∞, such that the first interval begins at time 0 and extends to time *s*_1_, and the last interval contains time up to infinity to avoid any orphaned events. In the ith interval, the birth rate is λ_*i*_, the death rate is *μ_i_*, the serial sampling rate is *ϕ_i_*. At the boundaries of the interval, we allow for each lineage to give birth, die, or be sampled with probabilities Λ_*i*_, *M_i_*, and Φ_*i*_ respectively. The intervals are left-inclusive so that the process is in interval *i* if *s*_*i*−*i*_ ≤ *t* < *s_i_*. We assume that at any event time *s_i_*, at most one of {Λ_*i*_, *M_i_*, Φ_*i*_} is nonzero (that is, we forbid there to be multiple types of events at a single event time).

To compute *E*(*t*), we need to compute *E_i_*(*t*) for all intervals, where *E_i_*(*t*) depends on all *E_j_*(*t*) for *j* < *i*. In this setup, we can analytically solve the differential equation for *E* and obtain,

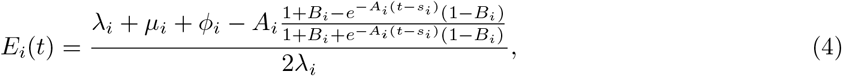

where

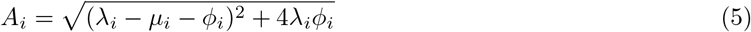

and

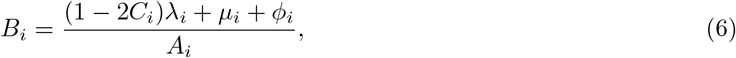

and where *C_i_* is defined as

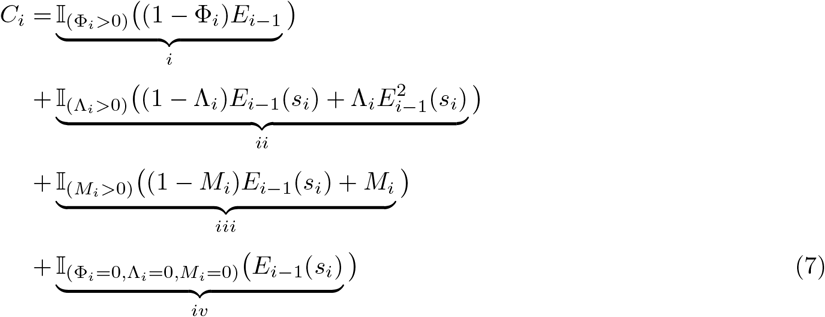

for *i* ≥ 1 and with *C*_0_ = 1. In *C_i_*, each element corresponds to one of the possible event types (or the absence of any event type), (i) this is an intensive sampling time where the lineage is unsampled, and the lineage is unobserved between time *s_i_* and the present, (ii) this is a burst time and either the lineage does not experience the burst event and is unobserved between time *s_i_* and the present, or it experiences the burst without having either descendant observed between time *s_i_* and the present, (iii) this is a mass extinction event and either the lineage survives the mass extinction but is unobserved between time *s_i_* and the present or the lineage dies in the mass extinction, (iv) there is no event at this time and the lineage is unobserved between time *s_i_* and the present. Terms (i)-(iv) are mutually exclusive as we forbid more than one event at any time.

### Probability along an observed lineage

Next, we define the differential equation computing the probability of a branch-segment along an observed lineage (Figure 5). A branch-segment terminates at a birth event or a sampling event, and over a small time Δ*t* we have,

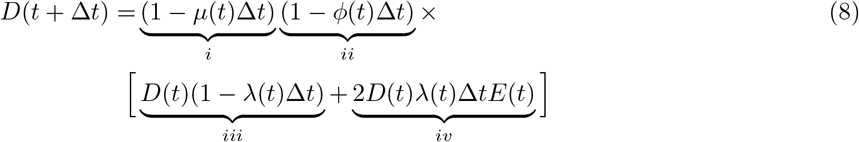

which accounts for the two scenarios that can apply over an interval, (1) no death (*i*) and no serial sampling (*ii*) and no birth (*iii*), (2) no death (*i*) and no serial sampling (*ii*) and a birth event (*iv*) but only one of the lineages is observed. As before, we can rearrange the above equation into a differential equation, yielding

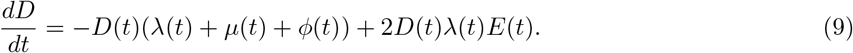

The analytical solution to this differential equation for piecewise-constant rates is

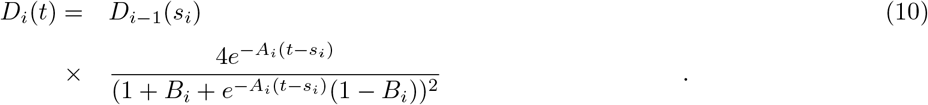

Note that this definition of *D*(*t*) includes only the continuous-time portions of the model, in other words we leave out the probabilities of all the events. This requires us to keep track of these separately, but makes our likelihood easier to compare to that of Gavryushkina et al. (Gavryushkina et al. 2014).

### Conditioning

When employing birth-death process models, we may wish to condition our model on having some tip samples (extinct or extant) in the tree or the tree surviving to the present day. Both options are dependent on whether we presume the tree begins at the origin (*t*_or_, with a single lineage) or at the most recent common ancestor (*t*_MRCA_, with two lineages). We provide a full description of the different conditioning options in the Supplementary Material.

### Validation of Theory and Implementation

Our generalized episodic fossilized-birth-death process is implemented in the open source software `RevBayes` (Höhna et al. 2016). We performed simulation based calibration to validate our implementation. We additionally validated our implementation within our simulation study testing the power and false-discovery rate of mass extinction events as well as the bias in speciation, extinction and fossilization rates. We provide more details of our validation of the underlying theory and implementation in the Supplementary Material.

### Priors and parameterization

To account for background variation in the rates of speciation, extinction, and sampling, we apply horseshoe Markov random field (HSMRF) prior distributions (Magee et al. 2020), which have been shown to be able to both detect rapid shifts in speciation rates and to reject time-varying models in favor of effectively constant-rate models. In a HSMRF model, we must specify prior distributions on the initial rates λ_1_, *μ*_1_, and *ϕ*_1_. We follow May et al. (May et al. 2016) in taking an empirical Bayes approach, first estimating parameters of a constant-rate fossilized-birth-death model, then fitting Gamma distributions to the posterior distributions of λ, *μ*, and *ϕ* and choosing the prior distributions to have twice the variance of the posterior distributions.

In our analyses, we discretize time into 100 evenly spaced epochs of 2.435 million years (*s*_0_, *s*_1_, *s*_2_,… = 0, 2.435, 4.870,…). At every time greater than 0, we allow for the possibility of a mass extinction event. We use reversible-jump Markov chain Monte Carlo (Green 1995; Freyman and Höhna 2018) in order to infer whether there is evidence for a mass extinction at each of these times. To determine the prior probability *p_i_* of a mass extinction at each event time, we first choose a prior expectation on the number of mass extinction events, 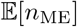, and set 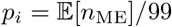. To account for prior sensitivity, we perform analyses with a range of prior expectations, 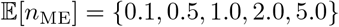. The mixtures are a *p*, (1 – *p*) mix of a mass extinction probability of 0 (no mass extinction), and a Beta(18,2) distribution (which has a prior mean of 0.9 for the extinction probability). To determine whether our results are robust to phylogenetic uncertainty, we also perform these 5 analyses on 6 distinct trees inferred by Wilberg et al. (Wilberg et al. 2019), for a total of 30 analyses.

### Convergence diagnostics

For all analyses (including those of simulated data), we ran 4 independent replicate MCMC simulations. To avoid convergence-related issues, we keep only the converged subset of these chains that pass a potential scale reduction factor (PSRF) cutoff (Brooks and Gelman 1998). As heavy-tailed distributions as the horseshoe distribution can cause problems for standard convergence diagnostics, we use the rank-stabilized PSRF of (Vehtari et al. 2020). After applying these standards (requiring PSRF < 1.01), we retained at least 3 chains for each empirical replicate analysis.

### Simulation study to test power and false-discovery rate

To determine the power of our method to detect mass extinctions, and its false positive rate, we performed analyses on 400 simulated trees. Of these, 200 are simulated from the posterior distribution on **λ**, ***μ, ϕ***, and ***M*** for tree T1 with a prior expectation of 0.5 mass extinctions (power analysis). The other 200 trees are simulated from a similar analysis but where no mass extinctions were allowed (false-positives analysis). By estimating diversification rates for a phylogeny that experience a mass extinction and mis-specifying the model by disallowing mass extinctions in the analysis, we allow for the possibility that the inferred diversification rates produce trees that appear to have experienced mass extinctions, providing a robust test of the false positive rate. The inference settings for these simulated datasets was identical to the empirical analyses.

### Code and data availability

The episodic fossilized-birth-death model is implemented in the software `RevBayes` (Höhna et al. 2016), available at https://RevBayes.github.io. Data and scripts for analyses and simulations are available at https://github.com/afmagee/gefbd. A tutorial about estimating diversification rates from phylogenies with extant and extinct taxa is available at https://revbayes.github.io/tutorials/divrate/efbdp_me.html.

## Acknowledgements

This work was supported by the National Science Foundation Graduate Research Fellowship under Grant No. DGE-1762114 and an ARCS Foundation Fellowship awarded to AFM. This research was supported by the Deutsche Forschungsgemeinschaft (DFG) Emmy Noether-Program HO 6201/1-1 awarded to SH.

