## supplemental_material for "Impact of K-Pg Mass Extinction Event on Crocodylomorpha Inferred from Phylogeny of Extinct and Extant Taxa"

#### Supporting Information

Andrew F. Magee[1] and Sebastian Höhna[2,3]

[1]*Department of Biology, University of Washington, Seattle, WA, 98195, U.S.A.*
[2]*GeoBio-Center, Ludwig-Maximilians-Universität München, Richard-Wagner Straße 10, 80333 Munich, Germany*
[3]*Department of Earth and Environmental Sciences, Paleontology & Geobiology, Ludwig-Maximilians-Universität München, Richard-Wagner Straße 10, 80333 Munich, Germany*

### S1 Estimated diversification rates incorporating phylogenetic uncertainty and prior sensitivity

To account for phylogenetic uncertainty in diversification rate estimates, we estimate diversification rates and mass extinctions on a total of 6 distinct trees from Wilberg et al. (2019), which we refer to as T1 to T6. Simultaneously, we investigate the sensitivity of our estimate to the prior expectation on the number of mass extinctions. Thus, we analyze each tree with a prior expectation of $\mathbb{E}(n_{\text{ME}}) = \{0.1, 0.5, 1.0, 2.0, 5.0\}$ mass extinctions. This leads to a total of 30 empirical analyses, which produce largely congruent results. In Figure S1 we plot the estimated rates of speciation, extinction, and fossilization, which is summarized in Figure 2 of the main text.

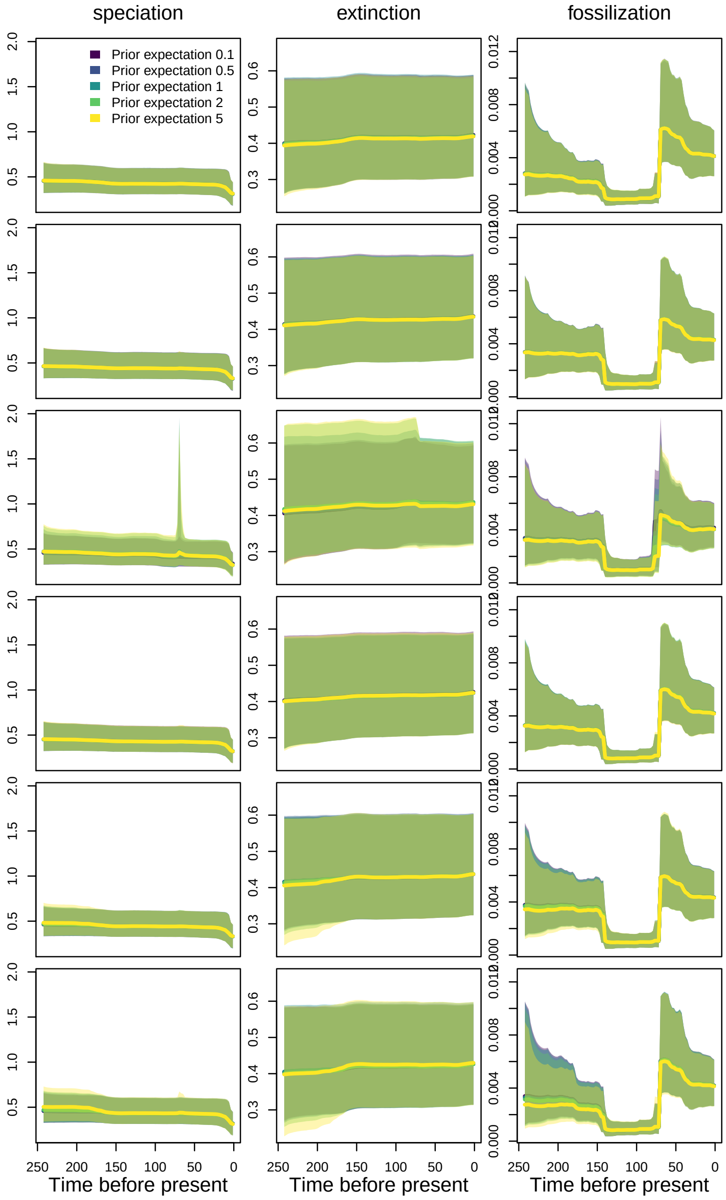

**Figure S1:** Estimated rates of speciation, extinction, and fossilization through time across all datasets and all priors on the expected numbers of mass extinctions. Datasets are shown in rows, while different priors on the expected number of mass extinctions are denoted by color. Solid lines are the posterior median rates, while shaded regions are the 90% CIs.

### S2 Additional empirical analyses

#### S2.1 Extinct and extant only phylogenies

We perform two analyses of subsampled trees to examine the contribution of fossil taxa to the signature of the mass extinction. First, we analyze a subtree of tree T1 consisting of only the extant Crocodylomorph taxa. This analysis detects no signal of the K-Pg mass extinction (Figure S3, top row), and the estimated speciation rate through time is effectively constant (Figure S2, top row). Both speciation and extinction rates are estimated to be lower than using the combined dataset (without fossils there is no fossilization rate to be estimated). Second, we analyze a tree consisting only of extinct Crocodylomorph taxa. This analysis strongly detects the K-Pg mass extinction (Figure S3, middle row). As with the extant-only analysis, the estimated speciation rate does not decrease towards the present, though it is otherwise similar to the diversification rate estimates obtained from the combined dataset (Figure S2). The extinction and fossilization rates estimated are almost identical to the combined analysis. Thus, at least for the Crocodylomorphs, the fossils provide the primary signal of the K-Pg mass extinction. Nevertheless, there is no harm in using a combined dataset. Diversification rates, however, do appear more sensitive to the exclusion of any taxa. Specifically, without the combined dataset it would not be possible to obtain the complete picture of historical diversification rates which includes the more ancient mass extinctions and recent decrease in speciation rate.

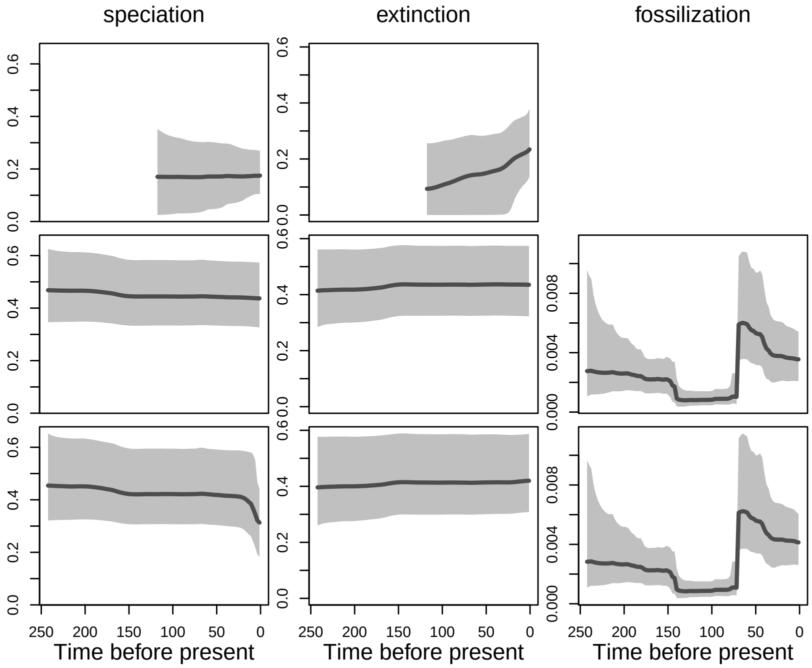

**Figure S2:** Estimated rates of speciation, extinction, and fossilization through time. Top row: only extant taxa used in analysis (hence no fossilization rate). Middle row: only extinct taxa used in analysis. Bottom row: all taxa used in analysis (reproduced from our main empirical analysis).

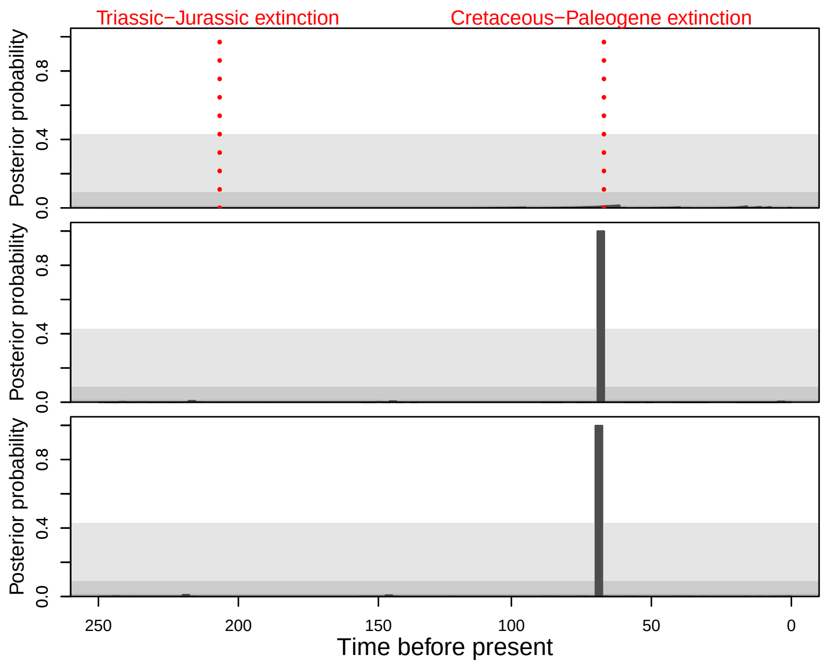

**Figure S3:** Support for mass extinctions. Top row: only extant taxa use in analysis. Middle row: only extinct taxa used in analysis. Bottom row: all taxa used in analysis (reproduced from our main empirical analysis).

#### S2.2 Assuming all fossils to be tips (treatment)

We used the phylodynamic treatment parameter to investigate the effect of assuming that all fossils are tips and not sampled ancestors. Specifically, we re-analyze the tree with $r_1 = r_2 = \cdots = r_{100} = 1.0$, which forces all tips to be fossils. This is not biologically meaningful, as leaving a fossil does not enforce the species to go extinct (there is no treatment), but this analysis provides insight into the effects and systematic bias of forcing fossils to all be tips.

We find that the estimated signal of the K-Pg mass extinction is robust to assuming that all fossil taxa must be tips (Figure S5). However, the estimated diversification rates are noticeably different (Figure S4). Estimated speciation and extinction rates are much lower (by a factor of two), while the fossilization rate is overall higher (by a factor of three). The speciation rate does not display any decrease towards the present, and the extinction rate increases towards the present day. Note however, in cases with $r > 0$ that fossilization also implies the death of a lineage, and the total death rate is actually $\mu(t) + \phi(t)r(t)$, which explains why the estimate extinction rate is lower when assuming that all fossils are tips.

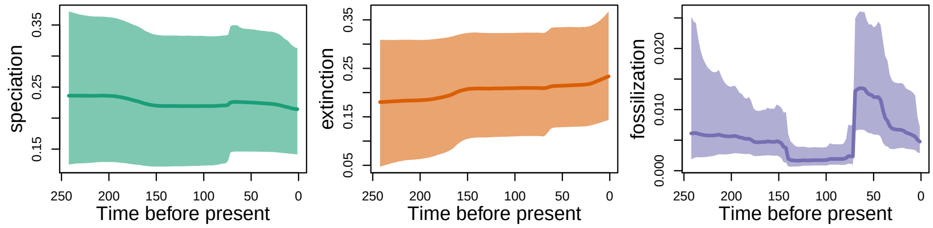

**Figure S4:** Estimated rates of speciation, extinction, and fossilization through time when assuming that all fossils are tips (treatment).

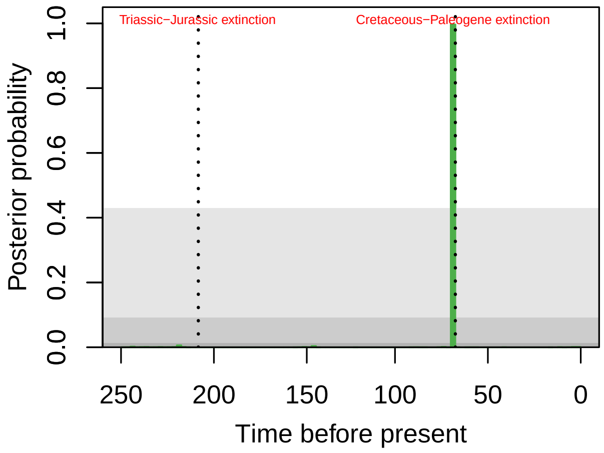

**Figure S5:** Support for mass extinctions at all 99 timepoints when assuming that all fossils are tips (treatment).

### S3 Simulated data analyses

We performed a simulation study to explore the power of our crocodylomorph analysis and the false positive rate. To assess power, we simulated trees from the posterior distribution of tree T1 with a prior expected number of 0.5 mass extinctions. We ensured that all simulated datasets experienced a mass extinction at the K-Pg boundary. To do this, we set the mass extinction death probability at the K-Pg to be 0.9 for any posterior sample that had a mass extinction death probability of less than 0.5 (this affects a very small proportion of simulations, as the estimated posterior probability of the K-Pg mass extinction was 0.998).

To assess any tendency for false positives, we fit diversification rates through time for the same dataset but without the possibility of mass extinctions. Disallowing mass extinctions in the real-data inference could lead to inferred rates that produce temporal signatures that look like mass extinctions (Crisp and Cook, 2009; Stadler, 2011; May et al., 2016). Trees simulated using parameter values drawn from the posterior distribution could appear to have mass extinctions and inference of simulated datasets may favor mass extinctions when there were none. Thus, this should provide a worst-case scenario on false positives. To ensure that we had sufficient resolution, we analyzed 250 trees for each scenario, and took the first 200 analyses that passed convergence cutoffs.

In the main text, we focused on the number of inferred mass extinctions per-dataset. In doing so, we used a 2 log Bayes factor threshold of 10 to determine if a mass extinction was detected or not. This cutoff is motivated by examining the distribution of all posterior probabilities in support of mass extinctions pooled across all break times and all simulated analyses. In the "false positive" analyses without mass extinctions, approximately 1% of all ($99 \times 200$) possible mass extinctions would be inferred to be significant at a Bayes factor cutoff of 2 (Figure S6), and there are a number of mass extinctions above a cutoff of 6. Thus, to cut down on spurious inference of mass extinctions, we use a cutoff of 10 for determining support for mass extinctions. The rationale behind the rather high significance threshold is multiple testing, because we tested jointly for 99 possible mass extinctions, one per epoch.

To assess the accuracy of our estimated rates through time, we use the mean relative absolute error, $1/n \sum_{i=1}^{n} [(\hat{\theta}_i - \theta_i)/\theta_i]$, where we take the posterior median as the parameter estimate. The distribution of relative errors for speciation and extinction are in general low (Figure S7). The accuracy for speciation and extinction rate estimates is comparable to what was found in Magee et al. (2020) in both their analyses where only speciation varied and their analyses where both speciation and extinction varied. The fossilization rate is apparently much more difficult to estimate, the average error is much higher (Figure S7). Further, where speciation and extinction rates are generally underestimated, the fossilization rate is generally overestimated. Estimation error is larger when compound parameters like net diversification ($\lambda(t) - \mu(t)$) are considered, suggesting that we are actually estimating the rates parameterized, and not compound parameters.

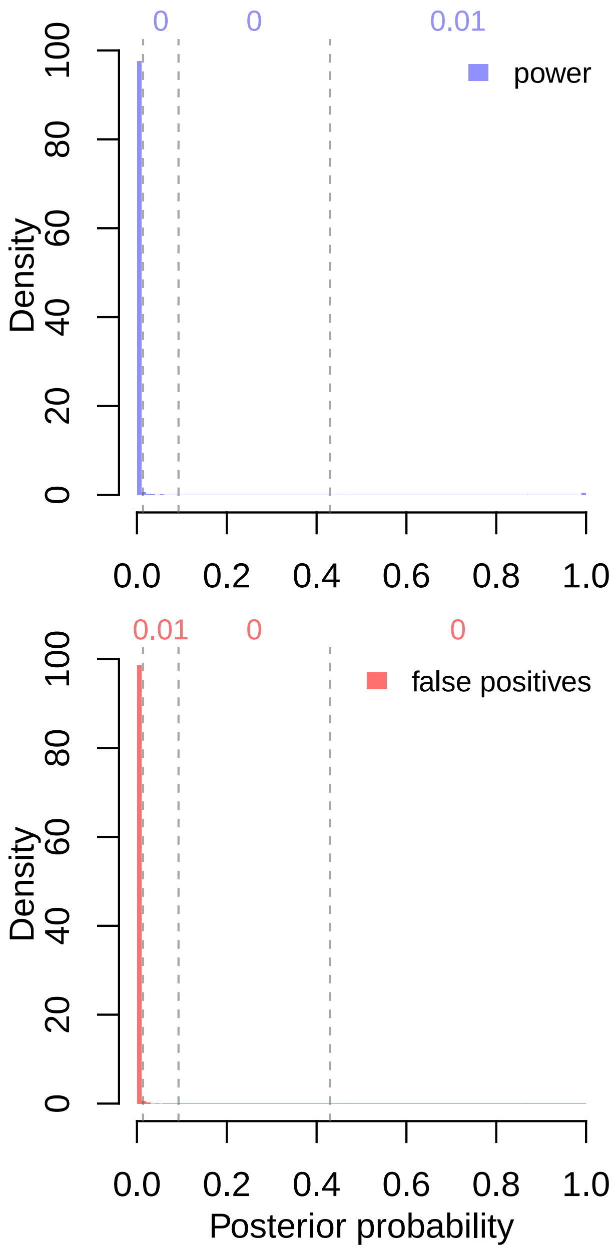

**Figure S6:** The posterior probability of a mass extinction in analyses of simulated data, pooled across all 99 times at which mass extinctions were allowed and all 200 simulated datasets. Vertical lines denote 2 log Bayes Factor cutoffs of 2, 6, and 10, which correspond to weak support, support, and strong support for a mass extinction. Numbers in each interval indicate the proportion of all posterior probabilities that fall in this interval, rounded to the nearest 1/100th.

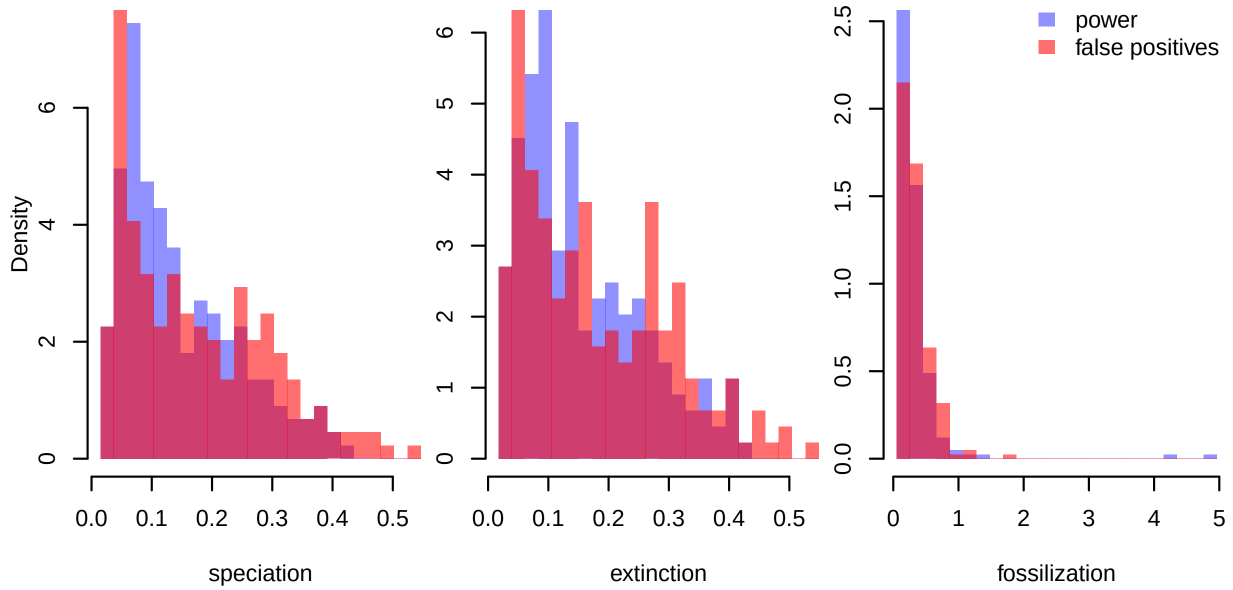

**Figure S7:** Accuracy of the estimated continuous parameters in the simulations. Accuracy is measured as the mean relative absolute deviation from the true rate, such that a value of 0.1 means an average absolute relative error of 10%.

### S4 Model adequacy

To assess the adequacy of our model performance, we employ posterior predictive simulations. Specifically, we simulate trees for the same mass extinction priors (0 and 0.5) and dataset (T1) as we use for our false positive and power analyses. For each of the three converged chains, we subsample to 2500 posterior samples, for each of which we simulate one tree, for a total of approximately 7500 trees (in a few cases the simulator failed). As mentioned above, in cases where the K-Pg mass extinction probability was estimated to be less than 0.5, we set the mass extinction probability to 0.9, this affects approximately 0.2% of the simulated datasets with mass extinctions. The first 250 of these trees include the same trees for which we performed our simulated data analyses.

We employ 16 summary measures of phylogenies, many of which are standard in the literature. They are:

1. Colless' (normalized) imbalance statistic, (Colless, 1982). Larger values mean trees are more imbalanced than expected under lineage-exchangeable models like the one derived in this paper.
2. Tree length, the sum of all branch lengths in the tree.
3. Tippyness, the proportion of all branch lengths that are edges subtending a (fossil or extant) tip (Fiala and Sokal, 1985; Rohlf et al., 1990).
4. Tippyness (extant), the proportion of all branch lengths that are edges subtending an extant tip. This statistic should be sensitive to misspecification of the random sampling model assumed for sampling events (Fiala and Sokal, 1985; Rohlf et al., 1990).
5. Tippyness (extinct), the proportion of all branch lengths that are edges subtending a fossil tip. Combined with tippyness (extant), this statistic allows one to localize issues with tippyness.
6. Longest branch, the length of the longest branch in the tree (Duchene et al., 2019). Computed such that sampled ancestors break up branches.
7. The gamma statistic of (Pybus and Harvey, 2000), a measure of the concentration of branch lengths towards the root of the tree.

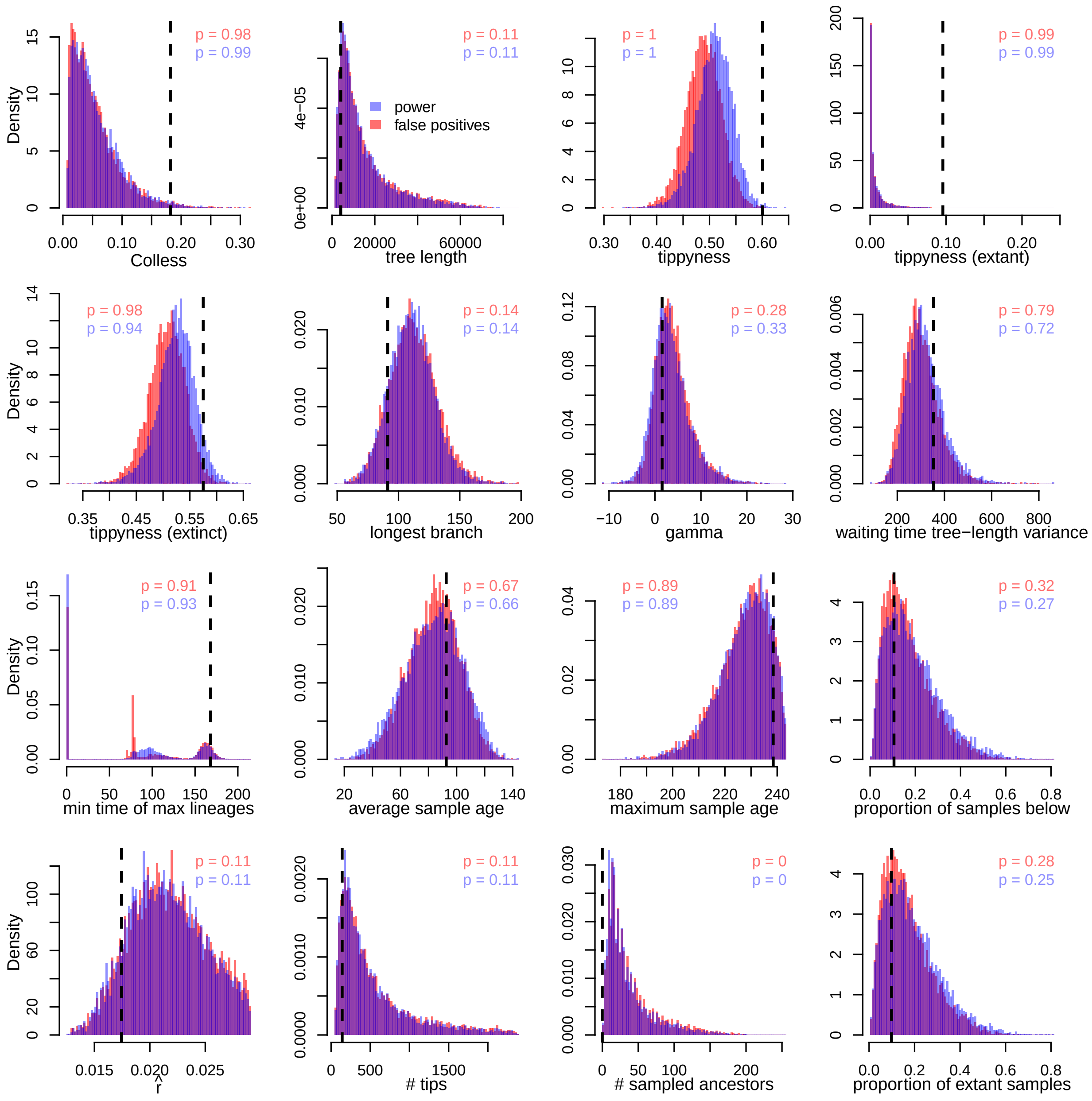

**Figure S8:** Posterior predictive distributions (red and blue) for each of 16 summary statistics. The observed value is shown in black. Posterior predictive p-values are rounded to the nearest 1/100th and are represent proportion of posterior predictive values below the observed value.

8. Waiting time tree-length variance, a measure designed to detect heterogeneity through time in the birth-death process. To compute, break the tree into intervals at every birth and sampling event. Let $\tau_i = n_i \Delta_i$ be the total tree length in time interval $i$, equal to the number of lineages in that interval multiplied by its duration in time. The statistic is then $\text{Var}(\boldsymbol{\tau})$.
9. Minimum time of maximum lineages, the most recent time in the lineage-through-time curve which has the maximum number of lineages, (Duchene et al., 2019). The minimum ensures uniqueness over multiple modes, though it means that for trees lacking serial samples the value will always be 0.
10. Average sample age, the average age of all samples (including extant tips, fossil tips, and sampled ancestors) (Duchene et al., 2019).
11. Maximum sample age, the oldest age of all samples (including extant tips, fossil tips, and sampled ancestors).
12. Proportion of samples below the youngest branching time. This measure should be sensitive to mis-estimating $\phi(t)$ relative to $\lambda(t) - \mu(t)$ in the recent past.
13. $\hat{r}$, a crude methods-of-moments estimator of the net diversification rate, (Magallon and Sanderson, 2001; Magee et al., 2020). This measure should capture whether the number of birth events in the trees are reasonable, relative to its age.
14. The number of tips in the tree, including extant and fossil tips.
15. The number of sampled ancestors in the tree.
16. The proportion of samples which are extant samples. When there is only event-sampling at the present ($\Phi_i = 0$ for $i > 0$), this statistic should be sensitive to how well $\phi(t)$ and $\Phi_0$ are matched.

Overall, we find that model performance is adequate. Few statistics exhibit very small or very large posterior predictive p-values. Furthermore, for a number of statistics, the mode of the posterior predictive distribution and the observed value appear to align, indicating good fit with respect to those statistics. However, the observed phylogeny has no sampled ancestors, while almost every simulated tree contains at least one sampled ancestor. This is likely at least in part an artifact of the tree building process assuming all samples are tips, which can be modeled by setting the phylodynamic recovery parameter $r$ to 1 (see above).

Colless' imbalance provides evidence for unmodeled among-lineage variation in diversification rates, as the observed tree imbalance is larger than most of the posterior predictive tree imbalances. The tippyness family of measures indicate some issues with the sampling model. Overall, predicted tippyness is lower than the observed tippyness. Looking at tippyness restricted to both extant and fossil tips, we can see that the larger driver here is the length of branches leading to extant tips. This could be explained if the 14 extant crocodylomorphs in the tree represent a diversified sample rather than a random one (Höhna et al., 2011). The tippyness restricted to fossil tips shows less misspecification, though there is still some discrepancy between the predicted and observed values. The lack of sampled ancestors could play a role here; fossil tips must have a branch subtending them, and this means that a tree with only fossil tips and no sampled ancestors will be longer than one where some fossil samples are sampled ancestors.

### S5 Estimated diversity through time

Once fit to the data, our models can be used to make inferences about the historical diversity through time of a group. These estimated diversity through time curves can be compared to other estimates, *e.g.,* from the fossil record, to further validate the results. However, note that we estimate species diversity through time and not the number of genera or families which is common in paleontological studies.

To estimate the diversity through time, for a set of posterior samples of diversification rates we simulate a complete tree, not allowing the tree to go extinct. Complete trees are necessary because reconstructed trees do not contain the record of all species alive at some time in the past, only those that contribute to the sample. Though since complete trees are quite large, this process is slow and running on a smaller number of posterior samples will most likely be needed. This procedure integrates out uncertainty in the diversification rates through time and appropriately explores the tails of the number of lineages in different intervals. In Figure S9, we show the inferred number of Crocodylomorph lineages through time using this approach (for the analysis of tree T1 with $\mathbb{E}(n_{\mathrm{ME}}) = 0.5$). Our model predicts a period of rapid growth up through the mid-to-late Jurassic, followed by a slow increase until the end Cretaceous mass extinction (K-Pg), which leaves a massive impact. Following the extinction, there is a period of slow growth into the Eocene, followed by a decline to the present.

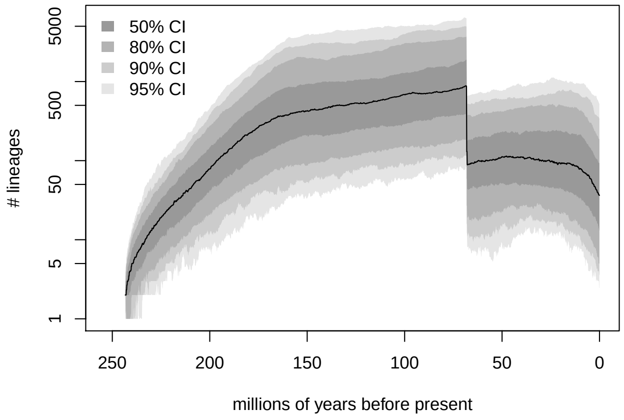

**Figure S9:** Posterior predictive distribution of the crocodylomorph diversity through time, median and 50% to 95% CIs. For each simulated tree, the number of lineages alive is binned into over 1000 intervals and recorded. All quantiles (median and CI) are taken per-interval.

### S6 Fossil tip ages

So far, we have shown that the signature of the K-Pg mass extinction is robust to (i) the choice of phylogeny (ii) the prior on the number of mass extinctions (iii) the exclusion of extant taxa (iv) the (tree inference) assumption that all taxa are tips. We have also shown that the model is largely adequate, and that any inadequacies are shared with models lacking mass extinctions. One factor that we have not addressed is the ages of the fossils. To assess robustness to fossil ages, we simulate 1000 trees based on T1 where we replace the ages with uniform draws from the stratigraphic ranges provided by Wilberg et al. (2019) (rejecting any draws of ages that would produce negative branch lengths). Fossil ages are likely important to identifying mass extinctions: simulated trees with mass extinctions often exhibit a band of fossil tips just prior to a mass extinction. By drawing new ages independently, we produce a sort of worst-case scenario where this signal gets maximally eroded.

Examination of the resampled LTT curves shows that there is still a clear drop (caused by the fossil tips), though there is uncertainty about the timing and magnitude of this drop (Figure S10). We can also compare summary statistics of these LTT curves to our posterior predictive distributions. Specifically, we can compare the number of fossils in the interval immediately prior to the observed K-Pg mass extinction to the predictive distributions from analyses with and without mass extinctions. For this comparison, we use the analysis with $\mathbb{E}(n_{\mathrm{ME}}) = 0.5$, and we normalize the number of fossils to the peak of the LTT curve for comparability between large trees and small trees (the model for mass extinctions kills a proportion of lineages, rather than a fixed number). The resampled LTT curves show a somewhat smaller drop than in the empirical tree T1, but the drop is more in line with trees simulated with mass extinctions than simulated without (Figure S11). Further, pooling both the interval immediately prior to the K-Pg with the next oldest recovers essentially the entire drop. Overall, these resampled datasets suggest that the signature of the K-Pg mass extinction is robust to the fossil ages.

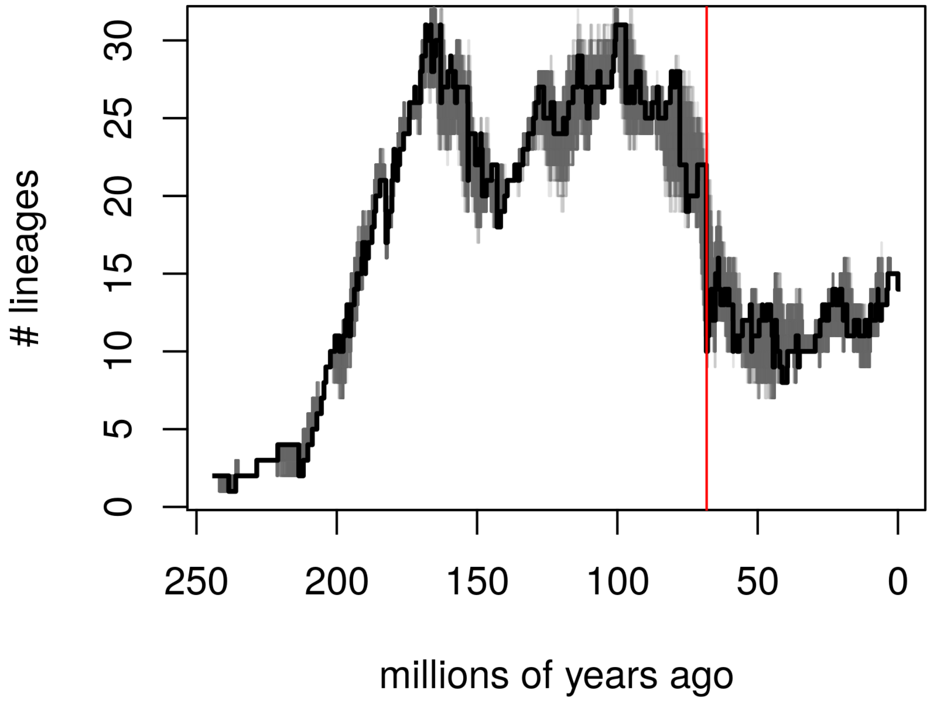

**Figure S10:** The LTT curve of T1 from Wilberg et al. (2019) (black), and 1000 LTT curves created by resampling the fossil times uniformly from the stratigraphic ranges. All resampled curves show a large drop around the time of the observed K-Pg mass extinction, suggesting they also contain evidence of the effect of the K-Pg.

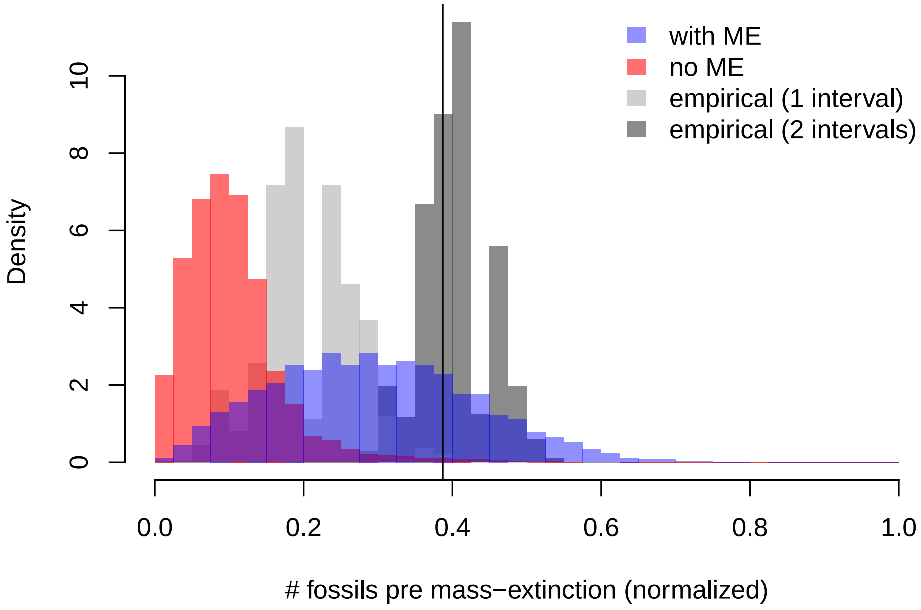

**Figure S11:** Distributions of the number of fossil samples in the time shortly before the observed K-Pg mass extinction for resampled versions of tree T1. For comparison between large and small trees, we normalize this number to the peak of the LTT curve. The blue histogram is the posterior predictive distribution based on the analysis with $\mathbb{E}(n_{\mathrm{ME}}) = 0.5$, while the red histogram is an analysis with no mass extinctions. The light grey histogram is the analog of the blue and red histograms, while the dark grey additionally includes fossils in the next oldest interval. The black line is the value in tree T1. Both resampled distributions show much larger numbers of fossil samples than expected without a mass extinction, suggesting that the signal of the K-Pg is robust to fossil times.

### S7 Interpretation of the terms in the Likelihood of the Generalized Episodic Fossilized Birth-Death Process

In the main text we only provided a brief explanation of our likelihood function. To make the explanation easier to understand, we reproduce the likelihood function again. The probability density of a phylogenetic tree $\Psi$ is

$$
\begin{aligned}
f(\Psi) = \quad & \frac{2^{I+H-||\mathcal{A}||-1}}{(I+H-||\mathcal{A}||)!} && (i)\\
& \times \prod_{t\in\mathcal{N}} \Big[\lambda(t)\Big] && (ii)\\
& \times \prod_{t\in\mathcal{F}} \Big[\phi(t)\big(r(t)+\big(1-r(t)\big)E(t)\big)\Big] && (iii)\\
& \times \prod_{t\in\mathcal{A}} \Big[\phi(t)\big(1-r(t)\big)\Big] && (iv)\\
& \times \prod_{i=1}^{l} \Big[\Lambda_i^{K_i}\big(2\Lambda_i E(s_i)+(1-\Lambda_i)\big)^{L(s_i)-K_i}\Big] && (v)\\
& \times \prod_{i=1}^{l} (1-M_i)^{L(s_i)} && (vi)\\
& \times \prod_{i=0}^{l} \Big[(1-\Phi_i)^{(L(s_i)-I_i)}\Phi_i^{I_i}(1-R_i)^{T_i}\\
& \qquad (R_i+(1-R_i)E(s_i))^{I_i-T_i}\Big] && (vii)\\
& \times \prod_{t\in\mathcal{B}} \Big[\frac{D(t_o)}{D(t_y)}\Big] && (viii)
\end{aligned}
\tag{S1}
$$

Term $(i)$ is the probability of the topology. There are $I+H-||\mathcal{A}||$ tips (fossil samples without sampled ancestors and extant samples) which have $(I+H-||\mathcal{A}||)!$ labelings. Furthermore, there are $(I+H-||\mathcal{A}||-1)$ internal nodes which have $2^{(I+H-||\mathcal{A}||-1)}$ left-right orientations. Since we do not consider left-right orientations in phylogenetics, the probability of the tree topology is $\frac{2^{I+H-||\mathcal{A}||-1}}{(I+H-||\mathcal{A}||)!}$.

Term $(ii)$ is the probability of the observed serial speciation in the tree (Figure 4g). Each of these happens with a probability density given by the speciation rate at that time.

Term $(iii)$ is the probability of the serially-sampled tips (Figure 4c-d). To be a tip, the sample must have no sampled descendants, which can occur in two ways. The sampling event may be treated, which happens with probability $\phi(t)r(t)$. Alternately, the sampled lineage may not be treated, and the lineage simply has no sampled descendants, which happens with probability $(\phi(t)(1-r(t))E(t))$.

Term $(iv)$ is the probability of the sampled ancestors (Figure 4c). We must sample the ancestor, and then it must go untreated (if it were treated, it would be a sampled tip).

Term $(v)$ is the probability of the observed and unobserved speciation events at tree-wide speciation burst events (Figure 4e). The probability of the observed burst speciation events is $\Lambda_i^{K_i}$. The probability of the lineages without observed burst speciation events is, $(2\Lambda_i E(s_i)+(1-\Lambda_i))^{L(s_i)-K_i}$. Lineages may not have observed burst speciation events for two reasons. A lineage might experience a burst speciation, but one of its children goes unsampled (leaving one continuous lineage in the reconstructed phylogeny), which happens with probability $2\Lambda_i E(s_i)$. Alternately, the lineage may not experience a burst speciation at all, which happens with probability $(1-\Lambda_i)$. In the case that there is no burst speciation at a particular interval time $s_i$ ($\Lambda_i = 0$) then there are no burst speciations ($K_i = 0$), and term $(v)$ is 1.

Term $(vi)$ is the probability of all lineages surviving a tree-wide mass extinctions events (Figure 4f). Each lineage that spans the $i$th mass extinction survives with probability $(1-M_i)$. We do not assume the possibility of observing any deaths at the time of the mass extinction.

Term ($vii$) is the probability of all the observed sampling times at given tree-wide sampling events (Figure 4g). This includes the probability of all the sampled lineages, $\Phi_i^{I_i}$, as well as the probability of all the unsampled lineages, $(1 - \Phi_i)^{(L(s_i)-I_i)}$. The probability of the sampled ancestors at this time is given by, $(1 - r(s_i))^{T_i}$, which is the probability that the sampled ancestors are not treated. The probability of the sampled tips is $(r(s_i) + (1 - r(s_i))E(s_i))^{I_i - T_i}$, which accounts for the possibility that the tip is treated $r(s_i)$, or that it is untreated but leaves no sampled descendants $(1 - r(s_i))E(s_i)$. In the case that there is no sampling event at a particular interval time $s_i$ ($\Phi_i = 0$) then there are no event samples ($I_i = 0$ and $T_i = 0$), and that term in the product collapses to 1.

Term ($viii$) is the probability of the observed branch segments (Figure 5). A branch segment is a portion of a branch that is uninterrupted by an interval time or an event (speciation, extinction, or sampling). The product of all branch segments yields the total probability of all the branches of the tree.

### S8 Different Conditions of the Generalized Episodic Fossilized Birth-Death Process

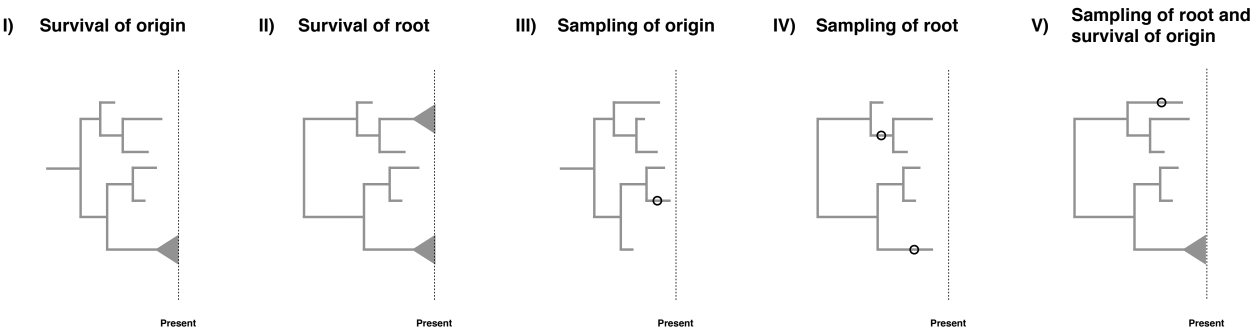

**Figure S12:** Five different possible conditions for our generalized fossilized-birth-death process. I) The process survives until the present. II) The process starts at the root and both descendants of the root survive until the present. III) Sampling at least one lineage. IV) The process start at the root and both descendants have at least one lineage sampled. V) The process starts at the root, both descendants have at least one lineage sampled, and the process survives until the present.

Birth-death models are often conditioned on specific events, see Stadler (2013) and Höhna (2015) for some discussion on the topic. However, when there are non-contemporaneous samples in the dataset which may be ancestral to other samples, conditioning becomes somewhat complex. The key issues for conditioning are whether it is assumed that the process starts at the root or the origin, and whether the descending lineage(s) is (are) assumed to leave any sampled descendant or specifically to have a descendant sampled at the present day. Consideration of these possibilities leads to five possible conditions, though conditioning is not strictly required.

**Survival of the origin** We condition the process on survival of one lineage, *i.e.,* at least on descendant of the lineage starting at the origin was sampled at the present. This condition represent a case when we have fossils and extant taxa and do not know if the fossils are stem fossils of the entire clade. The condition is obtained by computing $1 - E(t_{or})$ with $\phi(t) = 0$.

**Survival of the root** We condition the process on survival of both lineages, *i.e.,* at least one descendant of each lineage starting at the root was sampled at the present. This is the case for most macroevolutionary analyses without any fossils or if the fossils are known to belong within the crown group of the extant taxa. The condition is obtained by computing $(1 - E(t_{MRCA}))^2$ with $\phi(t) = 0$.

**Sampling of origin** We condition the process to have at least one sample being a descendant of the origin. This is simply a minimal condition that at least something was observed/sampled. This condition represents the case if we would also consider complete extinct clades. The condition is obtained by computing $1 - E(t_{or})$.

**Sampling of the root** We condition the process to require that both lineages starting at the root are sampled. In this case, all taxa might be extinct but the root age is known or inferred as a parameter of the model. The condition is obtained by computing $(1 - E(t_{MRCA}))^2$.

**Sampling of the root and survival of the origin** We condition the process on sampling of both descendant lineages of the root and at least one sample at the present. In this case, we condition on this specific root age but one of the descendant lineages of the root might have gone extinct while the other descendant lineage from the root must have survived. The condition is obtained by computing $(1 - E_{\phi(t)=0}(t_{MRCA}))(1 - E_{\phi(t)\neq 0}(t_{MRCA}))$.

For macroevolutionary analyses of diversification rates, condition (I) is the most adequate if we have both extinct and extant taxa, condition (II) if we have only extant taxa, and condition (III) if we have only extinct taxa. For phylodynamic applications, if it can safely be assumed that there are no sampled ancestors prior to the first observed infection (which will always be true if $r(t) = 1$), condition (IV) may be used, otherwise only condition (III) is applicable. Conditioning on survival as in (I), (II), or (V) requires $\Phi_0 > 0$, and so is primarily of interest in macroevolutionary applications. Of these conditions, (II) is the strictest and requires prior knowledge that the MRCA of the extant samples is the MRCA of all samples. Condition (V) is less restrictive, requiring only that none of the fossils could be sampled ancestors prior to the first observed speciation event, which would apply if all fossils are within the crown group. We could additionally condition on the number of extant taxa $N$, as suggest by Gernhard (2008), although there is, as of today and to our knowledge, no solution known to condition on the number of extinct taxa.

### S9 Comparison to the Gavryushkina Model

The model presented by Gavryushkina et al. (2014) represents the most parameter-rich model prior to the model in our paper, and is in fact a special case of this model. Specifically, the Gavryushkina model is the special case of ours where there are no mass extinction events ($M_i = 0 \ \forall \ i$), no birth bursts ($\Lambda_i = 0 \ \forall \ i$), and there is no distinction in treatment probability between $\phi$-sampling and $\Phi$-sampling ($R_i = r_i \ \forall \ i$). However, the presentation of the model in Gavryushkina et al. (2014) differs somewhat from ours, making a comparison between the two presentations useful.

For clarity, in our presentation of the Gavryushkina model, we keep our parameterization and notation. For readers looking at the original source material, our $\boldsymbol{\phi}$ is their $\boldsymbol{\psi}$ and our $\boldsymbol{\Phi}$ is their $\boldsymbol{\rho}$. Note also that there are some differences due to the choice of the direction of time. In our formulation, time for all terms flows backwards, thus $E_{i-1}(t)$ is an extinction probability for the interval preceding interval $i$. In the formulation of Gavryushkina et al. (2014), the preceding extinction probability would be $E_{i+1}(t)$ (or more accurately, $p_{i+1}(t)$).

#### S9.1 Terms A and B

Our term $\boldsymbol{A}$ is a generalization of that in the Gavryushkina model, in both cases we have,

$$A_i = \sqrt{(\lambda_i - \mu_i - \phi_i)^2 + 4\lambda_i\phi_i}. \tag{S2}$$

It can be seen that the term $\boldsymbol{B}$ in the Gavryushkina model is a special case of ours. Gavryushkina et al. (2014) define,

$$B_i = \frac{(1 - 2(1 - \Phi_i)E_{i-1})\lambda_i + \mu_i + \phi_i}{A_i}, \tag{S3}$$

and we define instead

$$B_i = \frac{(1 - 2C_i)\lambda_i + \mu_i + \phi_i}{A_i} \tag{S4}$$

where $C_i$ is defined as

$$\begin{aligned} C_i =& \mathbb{I}_{(\Phi_i>0)}\big((1 - \Phi_i)E_{i-1}\big) \\ &+ \mathbb{I}_{(\Lambda_i>0)}\big((1 - \Lambda_i)E_{i-1}(s_i) + \Lambda_i E_{i-1}^2(s_i)\big) \\ &+ \mathbb{I}_{(M_i>0)}\big((1 - M_i)E_{i-1}(s_i) + M_i\big) \\ &+ \mathbb{I}_{(\Phi_i=0,\Lambda_i=0,M_i=0)}\big(E_{i-1}(s_i)\big). \end{aligned} \tag{S5}$$

When there are no mass extinctions or birth bursts and $R_i = r_i$, this can be simplified to,

$$\begin{aligned} C_i =& \mathbb{I}_{(\Phi_i>0)}\big((1 - \Phi_i)E_{i-1}\big) + \mathbb{I}_{(\Phi_i=0)}\big(E_{i-1}(s_i)\big) \\ =& (1 - \Phi_i)E_{i-1}, \end{aligned} \tag{S6}$$

which is the same definition as in Gavryushkina et al. (2014).

#### S9.2 Extinction Probabilities

The extinction probability terms in Gavryushkina et al. (2014), $p_i(t)$, are a special case of our $E_i(t)$. Using our $s_i$ to represent the more recent boundary of the $i$th interval, Gavryushkina et al. (2014) define,

$$p_i(t) = \frac{\lambda_i + \mu_i + \phi_i - A_i \frac{e^{A_i(t-s_i)}(1+B_i)-(1-B_i)}{e^{A_i(t-s_i)}(1+B_i)+(1-B_i)}}{2\lambda_i}. \tag{S7}$$

We define

$$E_i(t) = \frac{\lambda_i + \mu_i + \phi_i - A_i \frac{(1+B_i)-e^{-A_i(t-s_i)}(1-B_i)}{(1+B_i)+e^{-A_i(t-s_i)}(1-B_i)}}{2\lambda_i}. \tag{S8}$$

When there are no mass extinction or birth burst events and $R_i = r_i$, $\boldsymbol{B}$ is the same in both models, and our definition is simply theirs where the last term on the numerator has been multiplied by

$$\frac{e^{A_i(t-s_i)}}{e^{A_i(t-s_i)}}.$$

Thus, $p_i(t)$ in Gavryushkina et al. (2014) is a special case of the $E_i(t)$ defined in this paper.

#### S9.3 Branch Probabilities

Despite the similarities in both terms $\boldsymbol{A}$ and $\boldsymbol{B}$, and the extinction probabilities, our $D_i(t)$ has no direct equivalent simpler case in the Gavryushkina et al. (2014) model. We define our $D_i(t)$ such that, for a branch that starts at time $t_o$ ends at time $t_y$ ($t_y < t_o$), the probability of observing that uninterrupted branch is $D(t_o)/D(t_y)$. Gavryushkina et al. (2014) define a similar quantity, $q_i(t)$,

$$q_i(t) = \frac{4e^{A_i(t-s_i)}}{(e^{A_i(t-s_i)}(1+B_i)+(1-B_i))^2}. \tag{S9}$$

We define $D_i(t)$ as

$$D_i(t) = D_{i-1}(s_i)\frac{4e^{-A_i(t-s_i)}}{((1+B_i)+e^{-A_i(t-s_i)}(1-B_i))^2}. \tag{S10}$$

In the simpler case where there are no mass extinctions or birth bursts and $R_i = r_i$, multiplying by

$$\left(\frac{e^{A_i(t-s_i)}}{e^{A_i(t-s_i)}}\right)^2,$$

shows us that

$$q_i(t) = \frac{D_i(t)}{D_{i-1}(s_i)}.$$

In essence, where $D_i(t)$ corresponds to the probability of an unbroken lineage between time $t$ and time 0, $q_i(t)$ track the probability of an unbroken lineage between time $t$ and the nearest younger interval time $s_i$. This difference is accounted for in Gavryushkina et al. (2014) by multiplying the likelihood by $q_{i-1}(t)^{L(s_i)-I_i}$ at every time $s_i$, where $L(s_i) - I_i$ is the number of lineages that are extant at the end of the interval, not counting the lineages sampled at the corresponding tree-wide event-sampling time.

### S10 Arranging terms in the likelihood

We note that our arrangement of terms in the likelihood is not the only possible option. We defined our branch segments such that they do not span multiple intervals and no birth bursts, intensive sampling events, or mass extinctions are possible. Because of this, our $D_i(t)$ reflect only the continuous rates $\lambda(t)$, $\mu(t)$, and $\phi(t)$, and the probabilities of birth bursts, intensive sampling events, and mass extinctions appear in separate (non-$D$) terms of the likelihood. We could instead have defined branch segments to only end at observed births and samples, in which case the branch segments would cross interval times and the probabilities of intensive sampling events, and mass extinctions would appear only in $D_i(t)$.

We also note that we can exploit the structure of the phylogeny to simplify the calculation for branch segments, term ($vii$) in the likelihood function. Along a single lineage, the probabilities of adjacent branch segments will cancel out because $t_y$ for one segment becomes $t_o$ for the next. For segments that begin with bifurcations, the addition of a new lineage means that a single $D(t_o)$ remains in the numerator. For segments that end in tips, there is no next segment, and thus $D(t_y)$ remains in the denominator. If we take $\mathcal{T}$ to be the set of all tip times, we can compute $(vii)$ as,

$$D(t_{or}) \prod_{t \in \mathcal{N}} D(t) \prod_{t \in \mathcal{T}} \frac{1}{D(t)}.$$

### S11 Related models

In Table S1, we present an overview of related work on birth-death processes. It should be noted that there are essentially two classes of papers in this list. There are papers that are concerned with the theory of computing the probability density of a tree given parameters, which often allow rates to be any time-varying function. There are also papers concerned with *inferring* diversification rates from phylogenies, which generally impose more restrictive assumptions on how rates may vary. Note that we are excluding here the literature on birth-death processes where the rate varies among lineages, which are beyond the scope of this paper. We also exclude comparison of models for sampling at the present, $\Phi_0$, a more thorough discussion of which is available in Höhna et al. (2011); Höhna (2014).

**Table S1: Time-varying birth-death models**
Parameters that are absent from a model are marked with a dash (–), and can be assumed to be 0 compared to a model that includes that parameter. Rates through time are classified by whether they are assumed to be constant (const.), piecewise constant or episodic (epis.), or whether they are allowed to be any time-varying function (any). Tree-wide events are either present (any) in a model or they are absent (–), except for tree-wide sampling which may be restricted to a single event at the present ($\Phi_0$). Conditioning includes conditioning on the various survival conditions discussed in Section S6 (I-V), and the number of tips (N). No conditioning listed is equivalent to simply conditioning on the time since the origin or MRCA. As many methods have been re-implemented in multiple software packages, the conditioning column only considers conditions used in likelihood equations in the cited paper. *Stadler (2010) and MacPherson et al. (2020) consider conditioning on the number of *extant* tips.

| Model and citation | $\lambda(t)$ | $\mu(t)$ | $\phi(t)$ | $r(t)$ | $\boldsymbol{\Lambda}$ | $\boldsymbol{M}$ | $\boldsymbol{\Phi}$ | $\boldsymbol{R}$ | Conditioning |
|---|---|---|---|---|---|---|---|---|---|
| Nee et al. (1994) | any | any | – | – | – | – | $\Phi_0$ | – | – |
| Gernhard (2008) | const. | const. | – | – | – | – | – | – | I, II, N |
| Stadler (2009) | const. | const. | – | – | – | – | $\Phi_0$ | – | I, II, N |
| Morlon et al. (2011) | any | any | – | – | – | – | $\Phi_0$ | – | II |
| Höhna (2014) | any | any | – | – | – | – | $\Phi_0$ | – | II, N |
| Stadler (2011) | epis. | epis. | – | – | – | any | $\Phi_0$ | – | II, N |
| Höhna (2015) | any | any | – | – | – | any | $\Phi_0$ | – | II, N |
| May et al. (2016) | epis. | epis. | – | – | – | any | $\Phi_0$ | – | II, N |
| Stadler (2010) | const. | const. | const. | 0 | – | – | – | – | II, III, N* |
| Stadler et al. (2012) | const. | const. | const. | const. | – | – | – | – | – |
| Stadler et al. (2013) | epis. | epis. | epis. | 1.0 | – | – | any | – | II |
| Gavryushkina et al. (2014) | epis. | epis. | epis. | epis. | – | – | any | – | II |
| (MacPherson et al., 2020) | any | any | any | ?? | – | any | any | – | I-V, N* |
| Present paper | epis. | epis. | epis. | epis. | any | any | any | any | I-V |
| Most general model | any | any | any | any | any | any | any | any | I-V, N |

### S12 Special Cases of the Birth-Death-Sampling-Treatment Process

In the following subsection we provide some special cases of our model. This simply shows that our model is a generalization of many previously published birth-death models, and how these models are related to another.

#### S12.1 Episodic birth-death process

We get the episodic birth-death process when we specify the parameters as follows:

$$\begin{aligned}\phi(t) &= 0\\ \Phi_i &= 0 \;\; \forall\ (i > 0)\\ M_i &= 0\\ \Lambda_i &= 0\end{aligned}$$

This simplifies our equations to

$$C_i = E_{i-1}$$

and

$$\begin{aligned}f(\Psi) = &\ \frac{2^{I-1}}{I!}\\ &\times \prod_{t\in\mathcal{N}} \Big[\lambda(t)\Big]\\ &\times \prod_{t\in\mathcal{B}} \Big[\frac{D(t_o)}{D(t_y)}\Big]\end{aligned}$$

#### S12.2 Episodic birth-death process with mass extinctions

We get the episodic birth-death process with mass-extinctions when we specify the parameters as follows:

$$\begin{aligned}\phi(t) &= 0\\ \Phi_i &= 0 \;\; \forall\ (i > 0)\\ \Lambda_i &= 0\end{aligned}$$

This simplifies our equations to

$$C_i = (1 - M_i)E_{i-1}(t_i) + M_i$$

and

$$\begin{aligned}f(\Psi) = &\ \frac{2^{I-1}}{I!}\\ &\times \prod_{t\in\mathcal{N}} \Big[\lambda(t)\Big]\\ &\times \prod_{i=1}^{l} (1 - M_i)^{L(s_i)} \qquad (vi)\\ &\times \prod_{t\in\mathcal{B}} \Big[\frac{D(t_o)}{D(t_y)}\Big]\end{aligned}$$

#### S12.3 Episodic fossilized-birth-death process

We get the (purely continuous) episodic fossilized-birth-death process for purely extinct taxa when we specify the parameters as follows:

$$\begin{aligned}r(t) &= 0\\ \Phi_i &= 0\\ M_i &= 0\\ \Lambda_i &= 0\end{aligned}$$

This simplifies our equations to

$$C_i = E_{i-1}(t_i)$$

and

$$\begin{aligned} f(\Psi) = & \frac{2^{H-||\mathcal{A}||-1}}{(H-||\mathcal{A}||)!} \\ & \times \prod_{t\in\mathcal{F}} \Big[(\phi(t)E(t))\Big] \\ & \times \prod_{t\in\mathcal{A}} \Big[\phi(t)\Big] \\ & \times \prod_{t\in\mathcal{N}} \Big[\lambda(t)\Big] \\ & \times \prod_{t\in\mathcal{B}} \Big[\frac{D(t_o)}{D(t_y)}\Big] \end{aligned}$$

#### S12.4 Skyline transmission process

We get the skyline transmission-process model (Gavryushkina et al., 2014) by specifying parameters as follows,

$$\Phi_i = 0$$
$$M_i = 0$$
$$\Lambda_i = 0$$

This simplifies our equations to

$$C_i = E_{i-1}(t_i)$$

and

$$\begin{aligned} f(\Psi) = & \frac{2^{H-1}}{H!} \\ & \times \prod_{t\in\mathcal{F}} \Big[\phi(t)\big(r(t) + \big(1-r(t)\big)E(t)\big)\Big] \\ & \times \prod_{t\in\mathcal{A}} \Big[\phi(t)(1-r(t))\Big] \\ & \times \prod_{t\in\mathcal{N}} \Big[\lambda(t)\Big] \\ & \times \prod_{t\in\mathcal{B}} \Big[\frac{D(t_o)}{D(t_y)}\Big] \end{aligned}$$

#### S12.5 Episodic transmission process with event-samples

We get the episodic sampled ancestor skyline model of (Gavryushkina et al., 2014) as follows,

$$M_i = 0$$
$$\Lambda_i = 0$$
$$R_i = r(s_i)$$

This simplifies our equations to

$$C_i = (1-\Phi_i)E_{i-1}(t_i)$$

and

$$
\begin{aligned}
f(\Psi) = \quad & \frac{2^{I+H-||\mathcal{A}||-1}}{(I+H-||\mathcal{A}||)!} \\
& \times \prod_{i=0}^{l} \Big[(1-\Phi_i)^{(L(s_i)-I_i)} \Phi_i^{I_i} (1-R_i)^{T_i} (R_i + (1-R_i)E(s_i))^{I_i - T_i}\Big] \\
& \times \prod_{t \in \mathcal{F}} \Big[\phi(t)\big(r(t) + \big(1-r(t)\big)E(t)\big)\Big] \\
& \times \prod_{t \in \mathcal{A}} \Big[\phi(t)(1-r(t))\Big] \\
& \times \prod_{t \in \mathcal{N}} \Big[\lambda(t)\Big] \\
& \times \prod_{t \in \mathcal{B}} \Big[\frac{D(t_o)}{D(t_y)}\Big]
\end{aligned}
$$

338 ## S12.6 Episodic transmission process with event-samples and perfect treatment

We get the "birth-death skyline" model of (Stadler et al., 2013) as a simplification of the episodic sampled ancestor skyline model by assuming perfect treatment as follows,

$$
\begin{aligned}
r(t) &= 1 \\
M_i &= 0 \\
\Lambda_i &= 0
\end{aligned}
$$

339 This simplifies our equations to

$$C_i = (1-\Phi_i)E_{i-1}(t_i)$$

340 and

$$
\begin{aligned}
f(\Psi) = \quad & \frac{2^{H+I-1}}{(H+I)!} \\
& \times \prod_{i=0}^{l} \Big[(1-\Phi_i)^{(L(s_i)-I_i)} \Phi_i^{I_i}\Big] \\
& \times \prod_{t \in \mathcal{F}} \Big[\phi(t)\Big] \\
& \times \prod_{t \in \mathcal{N}} \Big[\lambda(t)\Big] \\
& \times \prod_{t \in \mathcal{B}} \Big[\frac{D(t_o)}{D(t_y)}\Big]
\end{aligned}
$$

#### S13 Validation of likelihood function of episodic fosslized-birth-death process

The derivation of our likelihood function, *i.e.,* the probability density function of a phylogenetic tree, relies heavily on the extinction probability $E(t)$ and the probability of an observed lineage $D(t)$. In the main text we provided our mathematical derivations. Here, we additionally validate the analytical solutions using forward simulations under the generalized fosslized-birth-death process. We started the simulations with one single lineage and simulated forward in time starting at $T = \{0.01, 0.02, \ldots, 0.99, 1.0\}$ time units in the past. We chose $\lambda(t) = 1.0$, $\mu(t) = 0.9$ and $\phi(t) = 0.1$. Additionally, we divided the total time into four epochs, thus, allowing for tree-wide events at $t = \{0.25, 0.5, 0.75\}$ with probabilities $\Lambda = \{0.0, 0.0, 0.3\}$, $\mathbf{M} = \{0.0, 0.5, 0.0\}$ and $\Phi = \{0.2, 0.0, 0.0\}$. We repeated the simulations 100,000 times and recorded how often the process went extinct ($E(t)$) and how often exactly one lineage was observed ($D(t)$). Reassuringly, the probabilities obtained from the simulations and the analytical solutions match exactly (Figure S13).

The major novelty of our generalized birth-death-sampling process are the tree-wide events for mass extinctions and bursts of births. Thus, our simulations focusing on the three tree-wide events are sufficient as the impact of the continuous rates of speciation, extinction and sampling is validated through comparisons with the special cases in the previous section.

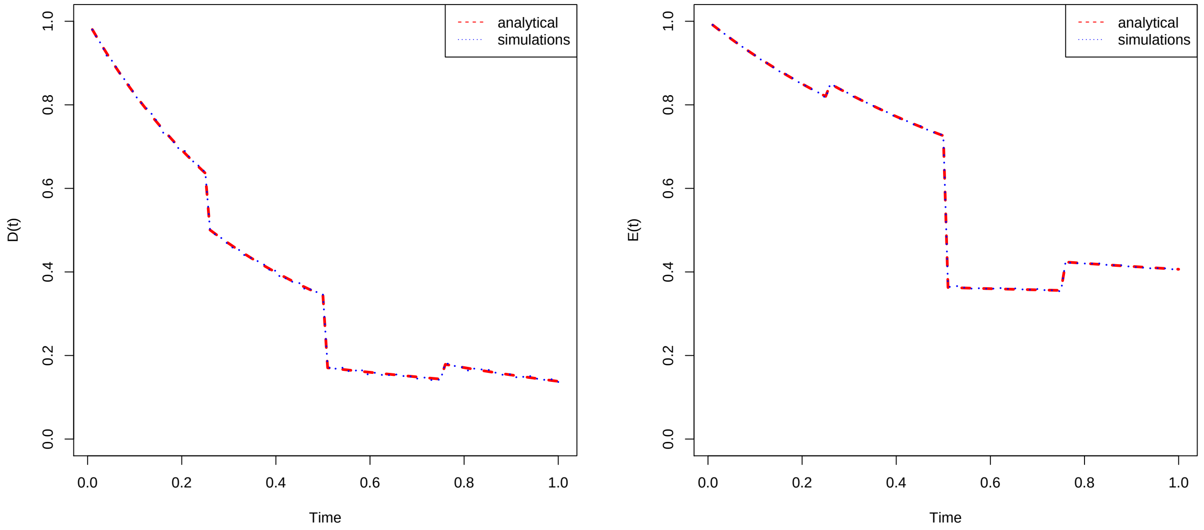

**Figure S13:** Comparing the analytical solutions for $E(t)$ and $D(t)$ with probabilities obtained by forward simulating the birth-death-sampling process. The analytical solutions match the expectations obtained through simulations.

### S14 Validation of likelihood function and implementation using simulation based calibration

We performed a simulation based calibration to validate our episodic fossilized-birth-death process. Standard theory of Bayesian inference defines that, if the data are generated under exactly the same model as used for inference, then the true parameter values are included in the credible intervals exactly with the frequency corresponding to the size of the credible interval (Huelsenbeck and Rannala, 2004; Cook et al., 2006). For example, the true parameter value should be covered in a 90% credible interval in 90% of the simulation replicates, neither more or less often. A nice feature of simulation based calibration is that the validation only works if all three, the simulation method, the likelihood function and the inference method (*e.g.,* the MCMC algorithm) are correctly implemented.

To validate our episodic fossilized-birth-death process, we choose the following approach. We designed a model with four equal-length epochs over a total time of 67.69 time units for the speciation, extinction and fossilization rates. Our assumption is that four epoch are sufficient to capture any potential problem with the per-epoch implementation but still being computationally manageable to perform thousands of MCMC analyses.

In principle, the choice of prior distribution does not matter. However, in practice, it is beneficial to choose realistic prior distributions so that trees simulated under parameters chosen from the prior distribution are reasonable, *i.e.,* are neither too large nor too improbable to survive. Thus, we specified a prior distribution on the net-diversification rate instead of the speciation rate to ensure that the simulated parameter values yield a positive net-diversification rate and hence the probability of the process going extinct is not close to 1.0. We employed a lognormal prior distribution on the net-diversification rate $\lambda_i - \mu_i$ with mean 0.01 and standard deviation 0.58, lognormal prior distribution on the extinction rate $\mu_i$ with mean 0.01 and standard deviation 0.58, and a lognormal prior distribution on the fossilization rate $\phi_i$ with mean 0.04 and standard deviation 0.58. Additionally, we employed a Beta$(20, 2)$ prior distribution on each the mass extinction death probability, the birth probability at a burst event, and the sampling probability at a tree-wide sampling event.

We implemented a forward simulator (which was also used for the posterior predictive distributions) and simulated trees given the parameter values drawn from the prior distribution. We conditioned the simulation on the root age of the extant tree (condition II, survival of the root). Then, we performed a standard MCMC algorithm using the same method as for the empirical analyses except that we used independent per-epoch priors instead of the HSMRF priors. The MCMC simulation was run for 10,000 iterations with 30 moves per iteration. We repeated this procedure 10,000 times to compute the frequency of how often the true parameter values were covered in the credible interval.

Finally, we computed and plotted the coverage frequencies for different credible interval sizes (Figure S14). The varying credible interval sizes help to validate that the posterior distributions are neither too peaked nor too flat due to heavy and light tailed distributions. Our results (Figure S14) demonstrate nicely that our forward simulator, our likelihood function, and our MCMC algorithm are all implemented correctly.

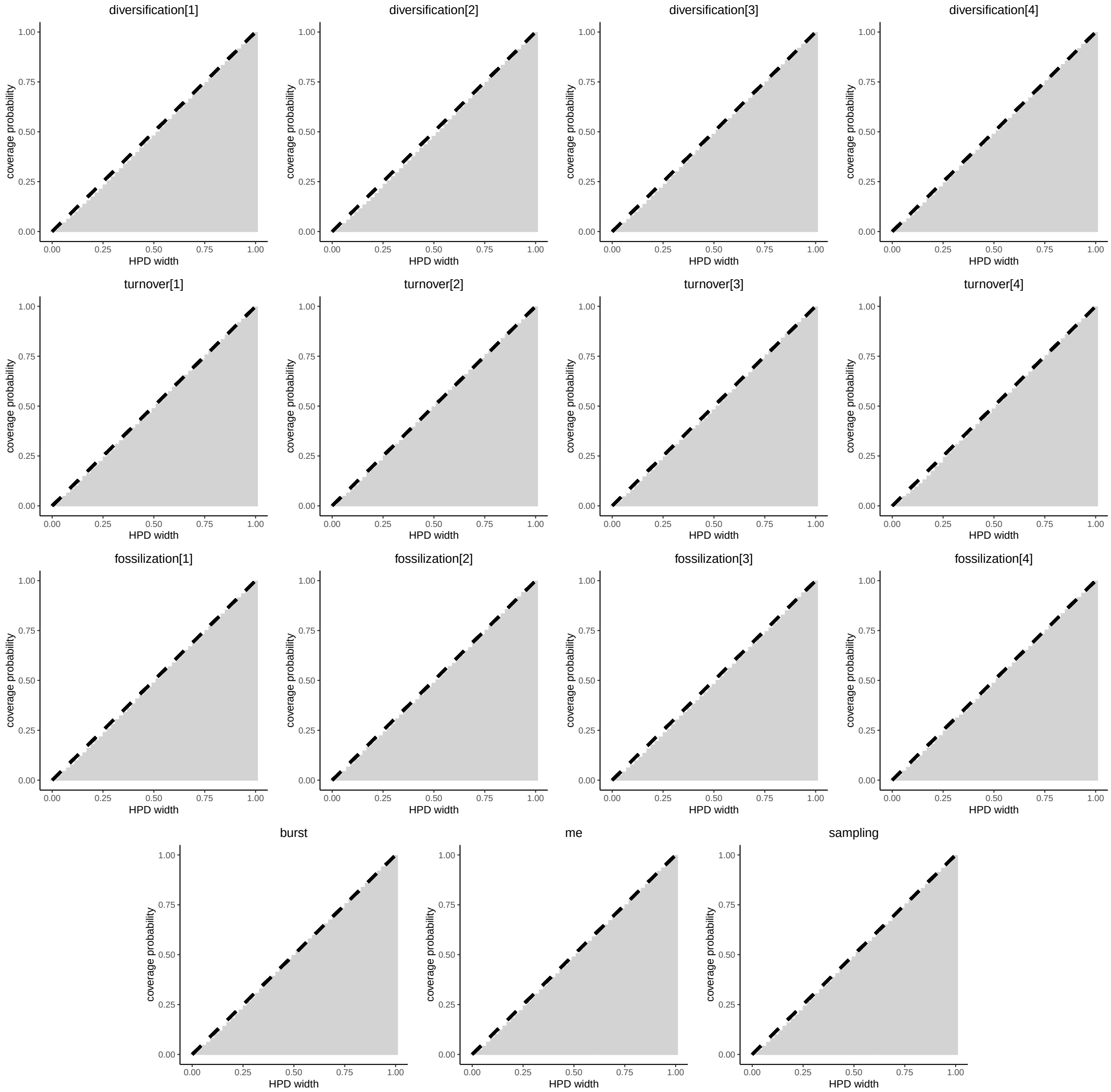

**Figure S14:** Validation of our derived likelihood function of the episodic fossilized-birth-death process with tree-wide events of burst of births, mass extinction, and sampling. We performed simulation based calibration and validated that the true parameter values are covered with the expected probability, *i.e.,* the size of the credible interval and the frequency of being including have to match. For all parameters in our example we observe a very good match between the expected and simulated coverage frequencies, indicating correct derivation of the theory and implementation of the likelihood function as well as MCMC algorithm.

### S15 Model parameterization

#### S15.1 HSMRF

In Listing 1 we provide the prior model specification for the speciation rates as employed in our analyses. `RevBayes` (Höhna et al., 2016) provides enormous flexibility in specifying how diversification rates vary through time and across lineages. While the model we have employed here has been shown to work well in certain circumstances (Magee et al., 2020), it remains open to the biologist and future work which type of diversification-rate variation is most prevalent and what model is most robust. Note that the speciation rate at present has two hyperparameters that are determined from a prior analysis of the dataset at hand using a constant-rate fossilized birth-death model.

```
speciation_at_present ~ dnGamma(speciation_rate_hyperprior_alpha,
                                 speciation_rate_hyperprior_beta)

speciation_global_scale ~ dnHalfCauchy(0,1)

for (i in 1:(NUM_INTERVALS-1)) {

  # Variable-scaled variances for hierarchical horseshoe
  sigma_speciation[i] ~ dnHalfCauchy(0,1)

  # non-centralized parameterization of horseshoe
  delta_log_speciation[i] ~ dnNormal( mean=0,
                                      sd=sigma_speciation[i]*
                                      speciation_global_scale*
                                      speciation_global_scale_hyperprior )
}

# Assemble first-order differences and speciation at present
#    into the random field prior for the speciation rate
speciation := fnassembleContinuousMRF(speciation_at_present,
                                      delta_log_speciation,
                                      initialValueIsLogScale=FALSE,
                                      order=1)
```

**Listing 1:** HSMRF on speciation rates.

We employed exactly the same type of model and priors on the extinction and fossilization rates. To keep this excerpt of our model concise, we show only the speciation rates.

#### S15.2 Improving MCMC

Applying the HSMRF prior distribution to birth, death, and fossilization rates can make MCMC difficult. We previously developed an MCMC framework for inference consisting of Metropolis-Hastings moves on the initial rate and a mixture of elliptical slice sampling and Gibbs sampling (Magee et al., 2020). This elliptical slice and Gibbs mixture works on the parameterization of the HSMRF prior in Listing 1, with the elliptical slice sampler working on the `delta_log_speciation` while the Gibbs sampler works on the `sigma_speciation` and `speciation_global_scale`. The Gibbs move as previously implemented updates all the `sigma_speciation` in order, then updates `speciation_global_scale`. As the `speciation_global_scale` parameter can be quite difficult to sample, we have implemented a move that is a $p$, $(1-p)$ mixture of the previous Gibbs update and a Gibbs update solely on `speciation_global_scale`. The conditionals involved in updating `speciation_global_scale` are unchanged, but as the `speciation_global_scale` parameter depends on both the vector `sigma_speciation` and the vector `delta_log_speciation`, more frequent updates to the `speciation_global_scale` parameter allow it to adjust more quickly to changes in `delta_log_speciation` (and vice-versa).

The other update is a simple swap move that operates jointly on the `delta_log_speciation` and the `sigma_speciation`. We outline this move in Listing 2; in brief, it simply swaps adjacent values of `delta_log_speciation[i]` and `sigma_speciation[i]` over the entire field. The move can migrate any pair (`delta_log_speciation[i]`, `sigma_speciation[i]`) to any pair (`delta_log_speciation[j]`, `sigma_speciation[j]`), however it does not add any *new* variation to the parameters, and thus the move can only be used to augment MCMC approaches that actually introduce new values into the vectors `delta_log_speciation` and `sigma_speciation`, such as the elliptical slice sampler and Gibbs mixture. The move is symmetric, and so the Hastings ratio is 0. The motivation for the move is as follows. The HSMRF prior enforces that most `delta_log_speciation[i]` are very small, such that the speciation rate contains a number of relatively flat regions interspersed with "jumps" where the rate changes more rapidly. In practice, there is often considerable uncertainty regarding exactly which intervals contain the jumps, and this move allows us to directly explore this uncertainty and move the jump locations around. Simultaneously, this move preserves the large-scale features of the speciation rate: for any pair of indices $i, j$ the total change in the speciation rate at $i$ and at $j$ will remain relatively consistent. The move operates on pairs of (`delta_log_speciation[i]`, `sigma_speciation[i]`) because these are compatible with each other; swapping a large-magnitude `delta_log_speciation[i]` with a small-magnitude `delta_log_speciation[j]` would pair a large-magnitude `delta_log_speciation[i]` with the small `sigma_speciation[j]` and this would lead to rejection.

```
u = randBernoulli(p=0.5)

start = floor(u)
end = start + 2 * (floor(length(delta_log_speciation) - start) / 2) - 1)

i = start

while (i < end) {
  tmp_d = delta_log_speciation[i]
  tmp_s = sigma_speciation[i]

  delta_log_speciation[i] = delta_log_speciation[i+1]
  sigma_speciation[i] = sigma_speciation[i+1]

  delta_log_speciation[i+1] = tmp_d
  sigma_speciation[i+1] = tmp_s

  i = i + 2
}
```

**Listing 2:** HSMRF swap move.
